# Glycolytic compensation rather than NAD⁺/NADH balance sustains neuronal function during mitochondrial stress

**DOI:** 10.64898/2026.08.20.746069

**Authors:** Shlesha Richhariya, Muskan Shah, Michael Rosbash

## Abstract

Neurons engage compensatory pathways that promote survival when confronted with mitochondrial dysfunction. Indeed, we recently showed that *Drosophila* neurons upregulate *Ldh* transcription to help survive the loss of the key mitochondrial fusion gene *Opa1.* Here, we further characterize this metabolic flexibility and show that it reflects a more general increase in glycolytic activity. A distinct mitochondrial perturbation, TFAM overexpression, similarly induces glycolytic gene expression including *Ldh* and also elevates lactate levels. LDH is also required to maintain neuronal function under *TFAM* overexpression. Notably, raising NAD^+^/NADH ratio by expressing the bacterial NADH oxidase LbNOX does not substitute for LDH function. On the contrary, it further compromises neuronal function in *Opa1*-deficient and *TFAM*-overexpressing neurons. Moreover, mitochondria-targeted LbNOX expression alone induces mitochondrial dysfunction and the compensatory glycolytic response. Together, these findings indicate that LDH-mediated rescue does not reflect an increase in NAD^+^/NADH ratio but is part of a broader neuroprotective metabolic reprogramming which enables neurons to withstand diverse forms of mitochondrial impairment.

## Introduction

The human brain consumes about 20% of total energy but comprises only about 2% of body weight. This reflects the unusually high metabolic demands of neurons (Magistretti and Allaman, 2015). Neurons are also special cells in other ways; they have unusual morphology and are post-mitotic as well as long-lived. Indeed, neurons often persist for the lifetime of an organism (Magrassi et al., 2013). It is therefore not surprising that disruptions in neuronal metabolism can contribute to age-related neurodegenerative disease (Camandola and Mattson, 2017; Murali Mahadevan et al., 2021). At the same time, neurons are quite resilient and can adapt their metabolism to energetic stress (Köhler-Solís and Schirmeier, 2025; Motori et al., 2020; Richhariya et al., 2025).

Mitochondria are central to cellular energy metabolism including that of neurons. Glycolysis converts glucose to pyruvate, which enters mitochondria and is converted to acetyl-CoA for oxidation in the TCA cycle; this process generates reducing equivalents such as NADH. NADH donates electrons to the electron transport chain (ETC), which lies within the inner mitochondrial membrane and drives ATP synthesis. Complexes I, III, and IV of the ETC use this electron transfer to pump protons, which creates the electrochemical gradient that drives the ATP synthase (Vercellino and Sazanov, 2022). Genetic or pharmacological disruption of electron transport chain function can therefore induce metabolic stress and impair neuronal function (Betarbet et al., 2000; Quintana et al., 2010).

A key feature of mitochondria is that they have their own small genome that contributes to their functions. In most animals including *Drosophila,* mitochondrial DNA encodes 13 essential subunits of the oxidative phosphorylation machinery, whereas the many other subunits and all other mitochondrial proteins are encoded by the nuclear genome (Rodrigues et al., 2022; Schmidt et al., 2010). Proteins imported into mitochondria include *Transcription Factor A, Mitochondrial* (*TFAM*). It is an essential mtDNA-binding protein, which is involved in mitochondrial genome packaging and maintenance. TFAM is conserved between mammals and *Drosophila* (Duncan et al., 2018; Larsson et al., 1998). Importantly, high levels of TFAM can disrupt mitochondrial function by excessively compacting mtDNA and altering mitochondrial gene expression (Bonekamp et al., 2021; Hunt et al., 2019).

Mitochondria also maintain function through a dynamic balance of fusion and fission. Fusion promotes functional complementation between individual mitochondria, whereas fission divides mitochondria and contributes to the removal of damage. In a previous study (Richhariya et al., 2025), we used tissue-specific CRISPR to disrupt mitochondrial fusion in *Drosophila* clock neurons. Despite severe mitochondrial fragmentation, knockout of the conserved inner-membrane fusion gene *Opa1* only caused a minor aging-dependent functional impairment. This was because these neurons mount a major transcriptomic response resembling the Warburg effect, in which several glycolysis-related genes were upregulated. Key among them was *Lactate dehydrogenase* (*Ldh*). Simultaneous loss of *Ldh* with *Opa1* led to much more severe impairment as well as age-related neurodegeneration. A similar but weaker transcriptional response was observed after loss of *Marf*, the conserved outer-membrane fusion protein, suggesting that this adaptation and *Ldh* upregulation is a general response to loss of fusion.

*Ldh* catalyzes the reaction of pyruvate to lactate while regenerating NAD^+^ from NADH; this sustains glycolysis and generates ATP (Gray et al., 2013). In highly glycolytic cancer cells, this NAD⁺-regenerating function of LDH can be essential for metabolic homeostasis and survival (Erdem et al., 2025). Thus, an important question arises: does the dependence on *Ldh* reflect a need for ATP via increased glycolytic activity, or a need for NAD⁺ regeneration for another purpose? NAD⁺ serves as a substrate for enzymes involved in cell signaling, chromatin regulation, and DNA repair beyond its role as a redox cofactor in glycolysis and other metabolic pathways. Moreover, the decline of NAD⁺ has been linked to aging and age-related disease (Covarrubias et al., 2021). In *Drosophila*, expression of the yeast NADH dehydrogenase *Ndi1* can extend lifespan (Bahadorani et al., 2010; Sanz et al., 2010). Ndi1 regenerates NAD⁺ by oxidizing NADH and transferring electrons to ubiquinone. Similarly, expression of LbNOX, which directly oxidizes NADH to NAD⁺ (Titov et al., 2016), extends lifespan but in a tissue- and sex-dependent manner (Yadav et al., 2026). These findings suggest that NAD⁺/NADH balance may influence age-related metabolic dysfunction and neurodegeneration.

Our previous study highlighted the importance of *Ldh* to fusion-deficient mitochondria, but it was uncertain whether this *Ldh*-dependence reflects a broad requirement for increased glycolytic activity or for NAD⁺ regeneration more generally, as described directly above. It was also uncertain whether the dependence extends to distinct forms of mitochondrial dysfunction beyond fusion. We address these issues here by testing how distinct forms of mitochondrial dysfunction interact with manipulations of compartment-specific NAD⁺/NADH balance, and by assaying whether enhancing NADH oxidation can replace the protective function of LDH. The results indicate that other forms of mitochondrial stress work similarly to the loss of fusion and that increased glycolytic activity, likely ATP generation, is the main purpose of the transcriptional upregulation including that of *Ldh*. These analyses also indicate that substantial NADH oxidation within mitochondria alone is sufficient to impair mitochondrial function and trigger compensatory glycolysis.

## Results

### Multiple glycolytic enzymes buffer neuronal vulnerability to *Opa1* loss

*Opa1*-deficient *Drosophila* clock neurons activate a transcriptional program that includes many glycolytic enzymes. They include *Ldh,* which is required to prevent neurodegeneration in the absence of *Opa1* (Fig. 1A; Richhariya et al., 2025). Elevated glycolysis together with increased *Ldh* expression should increase lactate production, and this expectation is consistent with a previous observation in *Drosophila* (Long et al., 2020). However, *Drosophila* LDH can catalyze the reversible interconversion of pyruvate and lactate (Price et al., 2025), making this prediction uncertain. We therefore measured lactate levels in clock neurons with cell type-specific CRISPR mediated loss of *Opa1* (*Opa1*-mut). The genetically encoded lactate sensor *CanlonicSF* (Aburto et al., 2022) in neurons indicated that lactate levels are indeed elevated in *Opa1*-mut *tdTomato*-labeled clock neurons as compared with *Cas9* controls (Fig. 1B).

**Figure 1:**
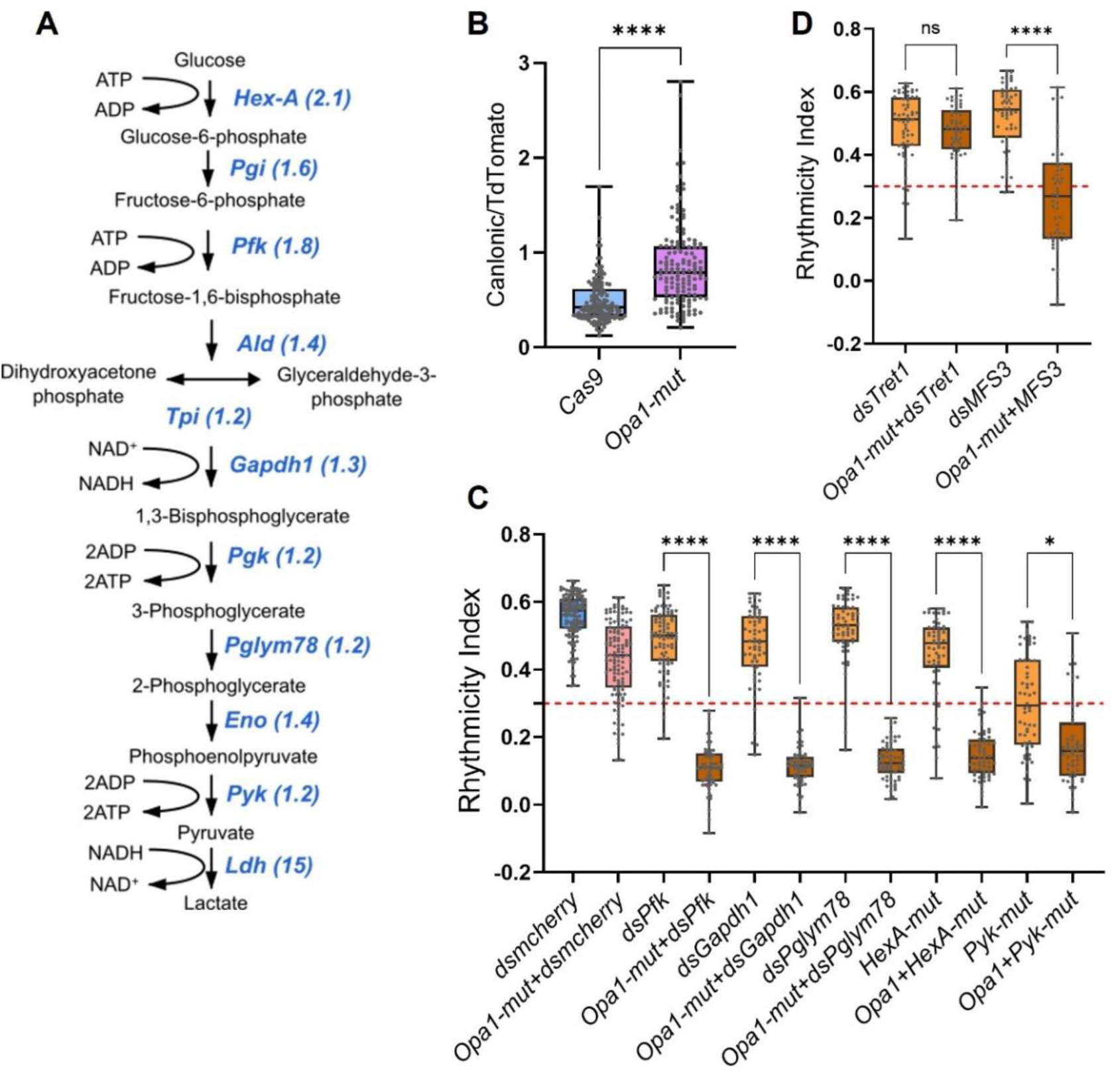
Glycolysis pathway genes compensate for loss of *Opa1* in clock neurons. **A.** Schematic of the glycolysis pathway with *Drosophila* enzymes shown in blue. Numbers in brackets indicate the fold upregulation in *Opa1-mut* clock neurons compared with *Cas9* controls (Richhariya et al., 2025). **B.** The ratio of *CanlonicSF* to *TdTomato* in lateral clock neurons from ∼3-week-old flies indicates higher lactate levels in *Opa1-mut* clock neurons. n>140 cell regions from ≥8 brains per genotype. ****p<0.0001, Mann-Whitney test. **C–D.** Rhythmicity index (RI), a measure of clock neuron function, for the indicated genotypes of ∼3-week-old flies. RI < 0.3, indicated by the red dotted line, is considered arrhythmic (for percent rhythmicity values, see Table 1). Several glycolytic or sugar transport genes, when knocked down using RNAi (*dsPfk, dsGapdh1, dsPglym78, dsMFS3*) or knocked out using cell-type-specific CRISPR (*Hex-A-mut, Pyk-mut*) in combination with *Opa1-mut* using *CLK856-Gal4; UAS-Cas9*, result in significantly lower rhythmicity than either perturbation alone, indicating that these genes compensate for *Opa1* loss and protect clock neuron function. RNAi against mcherry (*dsmcherry*) used as a control. *p<0.05; ****p<0.0001 for double perturbations compared with both single perturbations, Kruskal-Wallis test followed by Dunn’s post hoc test. For additional genes and details, see Table 1.

**Table 1:**
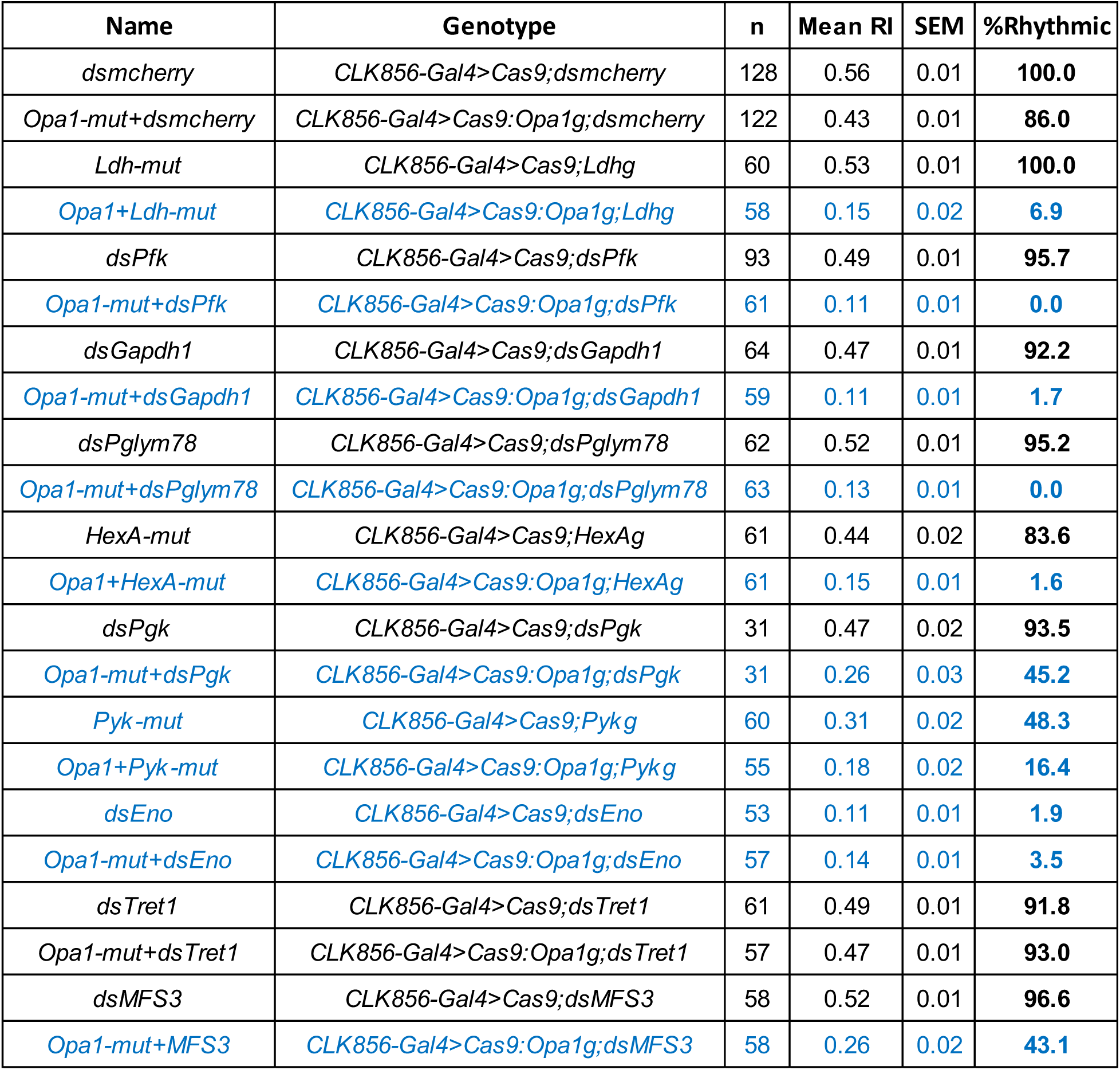
Interaction of glycolytic genes with *Opa1* in clock neurons. Mean rhythmicity index (RI), S.E.M., and percentage of rhythmic flies for the indicated genotypes from ∼3-week-old flies. Except for genotypes involving *Ldh, Pgk,* and *Eno*, the data are the same as those shown in Fig. 1C–D. Genotypes highlighted in blue are considered arrhythmic (mean RI < 0.3 and <50% rhythmic flies).

We then asked whether there is a broad requirement for glycolytic activity in the absence of *Opa1*. As done previously for *Ldh* (Richhariya et al., 2025) we knocked down several glycolytic genes using RNAi or knocked them out using cell-type-specific CRISPR (Fig. 1A, see Methods for details) in combination with *Opa1-*mut. Knockdown of *Pfk, Gapdh1, Pglym78*, *Pgk* or CRISPR-mediated knockout of *Hex-A (Hex-A-mut)* using the *CLK856-Gal4* driver alone had little or no effect on neuronal function as indicated by rhythmicity index or percentage of rhythmic flies (Fig. 1C; Table 1). This is consistent with previous work showing that *Drosophila* neurons can survive without glycolysis (Volkenhoff et al., 2015). In contrast, knockdown or knockout of each of these genes in combination with *Opa1*-mut caused an apparent complete loss of neuronal function, comparable to that observed with the *Opa1*-mut and *Ldh*-mut combination (Fig. 1C; Table 1).

Two glycolytic genes produced phenotypes even when knocked down alone. *Eno* knockdown had the strongest effect, whereas CRISPR mediated knockout of *Pyk* (*Pyk-mut*) resulted in a partial loss of function (Fig. 1C; Table 1). *Eno* and *Pyk* act late in glycolysis (Fig. 1A), suggesting that these steps may be particularly important for neuronal function. Consistent with the general picture, the partial phenotype caused by *Pyk* knockout was enhanced in combination with the *Opa1*-mut (Fig. 1C; Table 1).

Is increased sugar import important for preserving neuronal function? We used RNAi lines against the two sugar transporters with upregulated transcripts, *Tret-1* (Kanamori et al., 2010) and *MFS3* (McMullen et al., 2021). Knockdown of either transporter alone did not impair neuronal function (Fig. 1D; Table 1), suggesting that these transporters are not individually required under baseline conditions. Simultaneous knockdown of *Tret-1* with *Opa1*-mut also had no effect, but a similar double knockdown of *MFS3* resulted in reduced rhythmicity (Fig. 1D; Table 1) indicating that MFS3 is specifically required to maintain neuronal function during mitochondrial dysfunction. These results taken together indicate that mitochondrial dysfunction in *Opa1*-deficient neurons is accompanied by a functional requirement for glycolytic activity. This includes elevated sugar uptake and enhanced *Ldh* expression, which is also reflected by increased lactate levels.

### TFAM overexpression also induces LDH-dependent glycolytic compensation

Is the glycolytic response specific to loss of mitochondrial fusion through loss of *Opa1* or *Marf* (Richhariya et al., 2025), or does it reflect a broader response to mitochondrial dysfunction? To this end, we used two well-established perturbations of mitochondrial function: knockdown of the Complex I gene *ND-75* (Fig. S1A; Granat et al., 2023) and overexpression of the mitochondrial transcription factor *TFAM* (Fig. 2A; Cagin et al., 2015). Both caused the upregulation of several glycolytic genes in old clock neurons, and *Ldh* showed the strongest and most significant upregulation in both cases (Fig. 2B; Fig. S1B); this is similar to the *Opa1-*mut (Fig. 2B, D; S1B-C; Richhariya et al. 2025). The extent of glycolytic gene upregulation correlated with *TFAM* levels in *TFAM*-overexpressing clock neurons (*TFAM*-OE) (Fig. 2A, B, D). *ND-75* knockdown caused a mild effect on neuronal function, as seen by a small reduction in rhythmicity and a longer period. Loss of rhythmicity was further enhanced by the additional loss of *Ldh* (Fig. S1D, E). However, this interaction was milder than that between *Opa1* and *Ldh* (compare Fig. S1D to Table 1), suggesting that mitochondrial function may be less severely impacted by the *ND-75* knockdown than by *Opa1-*mut.

**Figure 2:**
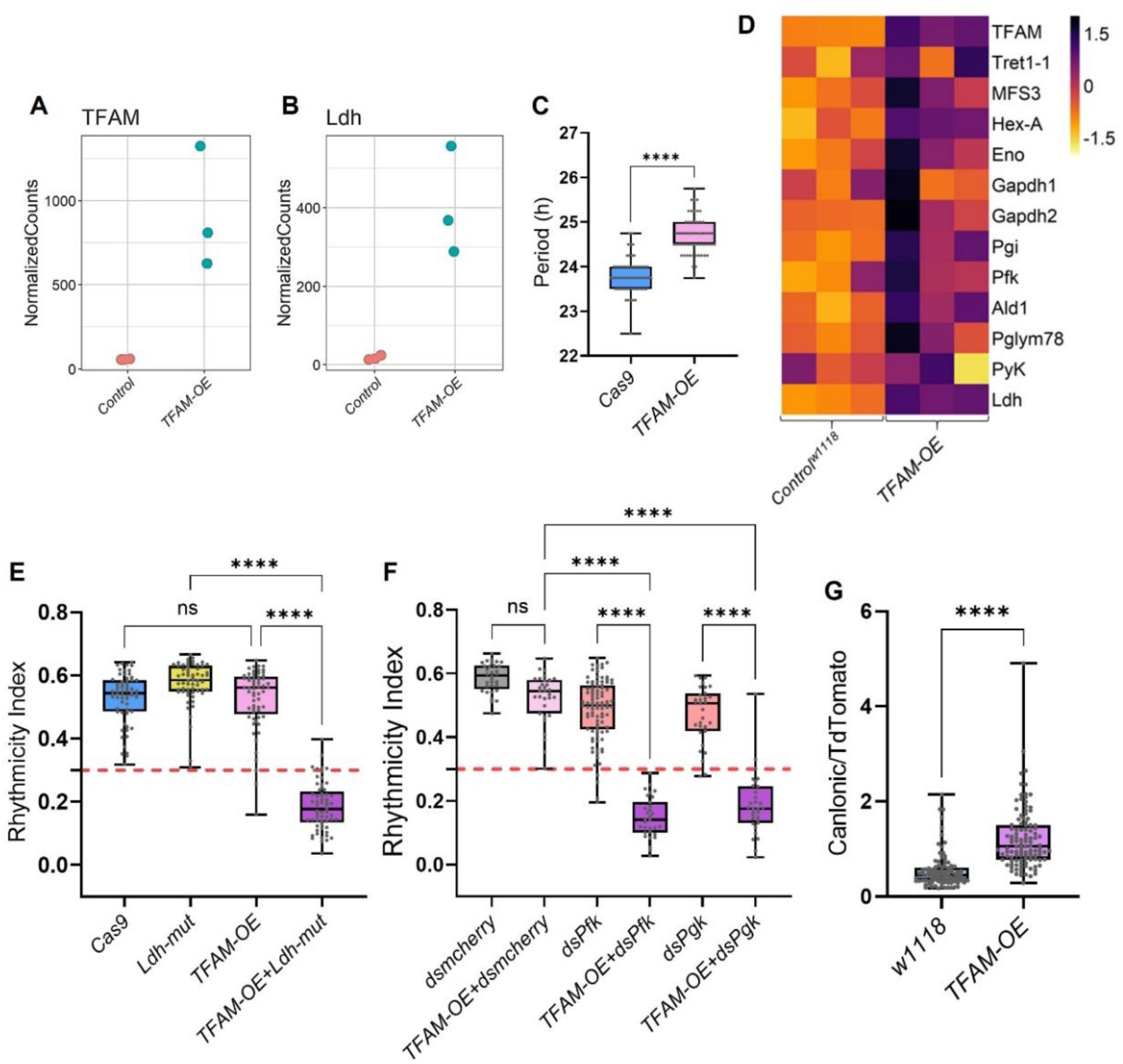
Upregulated glycolysis and *Ldh* compensate for decreased function in *TFAM*-overexpressing clock neurons. **A–B.** Normalized counts for *TFAM* (A) and *Ldh* (B) from bulk sequencing of ∼35-day-old control and *TFAM*-overexpressing (*TFAM-OE*) clock neurons, confirming *TFAM* overexpression and showing strong *Ldh* upregulation. **C.** Circadian period of rhythmic flies (RI > 0.3) shows a longer period in flies with *TFAM-OE* in clock neurons using *CLK856-Gal4* than in *Cas9* controls. **D**. Heatmap showing expression levels of genes involved in glycolysis and sugar transport across three replicates of indicated genotypes. Glycolytic genes and sugar transporters show a general increase in expression upon *TFAM-OE*; *TFAM* levels are shown in the top row. **E–F**. Rhythmicity index (RI), a measure of clock neuron function, for the indicated genotypes from ∼3-week-old flies. RI < 0.3, indicated by the red dotted line, is considered arrhythmic. Cell-type-specific CRISPR-based knockout of *Ldh* (*Ldh-mut*) or RNAi-based knockdown of glycolytic genes (*dsPfk, dsPgk*) in combination with *TFAM-OE* results in significantly lower rhythmicity than either perturbation alone. *dsPfk* and *dsPgk* data is the same as shown in Fig. 1C or Table 1. RNAi against *mCherry* (*dsmCherry*) was used as a control. n ≥ 30 flies per genotype. ****P < 0.0001, Kruskal–Wallis test followed by Dunn’s post hoc test. G. The ratio of *CanlonicSF* to *TdTomato* in lateral clock neurons from ∼3-week-old flies indicates higher lactate levels in *TFAM-OE* clock neurons compared with *w1118* controls. n>96 cell regions from ≥8 brains per genotype. ****P < 0.0001, Mann–Whitney test.

*TFAM*-OE in contrast was very similar to *Opa1*-mut (Richhariya et al., 2025). *TFAM*-OE alone had no effect on rhythmicity and only caused a small increase in period length (Fig. 2C, E). Similarly, *TFAM*-OE combined with loss of *Ldh* or with knockdown of the glycolytic genes *Pfk* and *Pgk* caused complete arrhythmicity (Fig. 2E,F); this is just like these combinations with *Opa1-mut*. Lactate levels were also similarly elevated in neurons with *TFAM*-OE (Fig. 2G). All these data indicate that *TFAM-OE* neurons rely on increased glycolytic activity and further underscore that different initiating mitochondrial perturbations drive a shared glycolytic response that helps preserve neuronal function.

### Compartment-specific NAD^+^/NADH manipulation does not substitute for LDH in neurons with impaired mitochondrial function

LDH converts pyruvate to lactate, consistent with the increased lactate levels observed in *Opa1*-mut and *TFAM*-OE clock neurons (Figs. 1B, 2G). However, this reaction also oxidizes NADH to NAD^+^, a key cofactor for many cellular processes. In proliferative cells, the role of LDH-mediated lactate production in regenerating NAD^+^ is considered to be more critical than its contribution to ATP production via glycolysis (Luengo et al., 2021; Titov et al., 2016). If this were the case in clock neurons, elevating the NAD^+^/NADH ratio in another way might replace the need for LDH under mitochondrial dysfunction conditions. To this end, we generated UAS lines expressing either LbNOX or the mitochondrially targeted mitoLbNOX, both of which oxidize NADH to regenerate NAD^+^ (Titov et al., 2016).

Expression of LbNOX alone did not detectably impact neuronal function, nor did its expression affect the arrhythmic phenotype of *Opa1+Ldh-*mut or *TFAM*-OE + *Ldh*-mut neurons (Fig. 3A-E). LbNOX did, however, marginally reduce the function of *Opa1*-mut and *TFAM*-OE clock neurons (Fig. 3B, D), suggesting that cytosolic NADH levels may help support LDH-dependent compensation. (See Discussion.)

**Figure 3:**
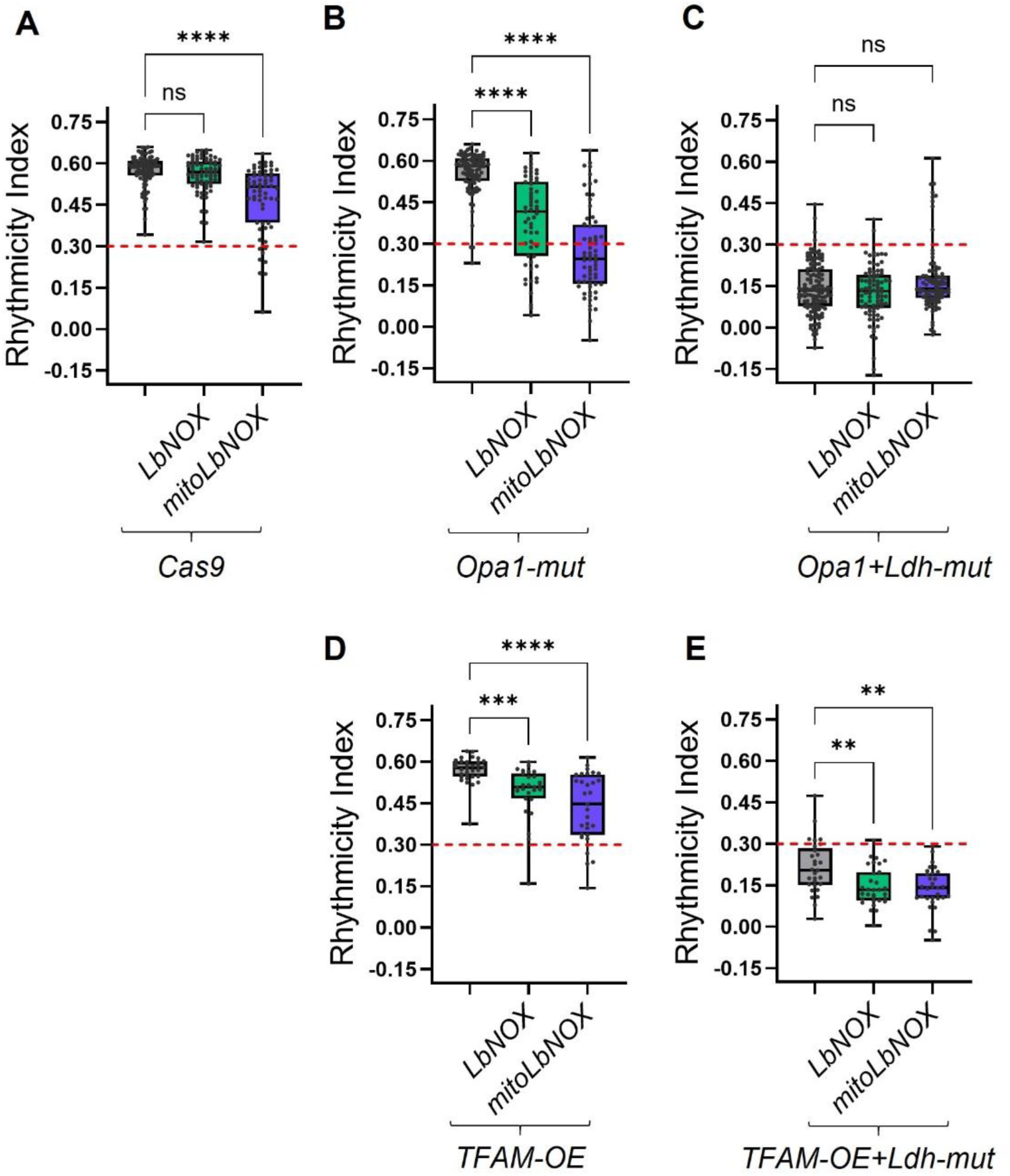
An elevated NAD+/NADH ratio cannot replace LDH function in clock neurons with mitochondrial dysfunction. **A–E.** Rhythmicity index (RI), a measure of clock neuron function, for the indicated genotypes driven by *CLK856-Gal4* from ∼3-week-old flies. RI < 0.3, indicated by the red dotted line, is considered arrhythmic. n ≥ 29 flies per genotype. **P < 0.01; ***P < 0.001; ****P < 0.0001; ns, not significant (P > 0.05), Kruskal–Wallis test followed by Dunn’s post hoc test. Expression of LbNOX in clock neurons does not alter rhythmicity, whereas mitoLbNOX causes a small but significant reduction in rhythmicity (A). Expression of either LbNOX or mitoLbNOX does not replace LDH function in *Opa1-mut* or *TFAM-OE* clock neurons (C, E). Expression of either LbNOX or mitoLbNOX modestly but significantly reduces rhythmicity in *Opa1-mut* and *TFAM-OE* clock neurons (B, D).

Mitochondria-targeted LbNOX (mitoLbNOX; Titov et al., 2016) is designed to have the same effect but specifically in mitochondria. Its expression alone caused a small reduction in rhythmicity and just like LbNOX did not replace LDH function in either *the Opa1*-mut or *TFAM*-OE backgrounds (Fig. 3A, C, E). Expression of mitoLbNOX also reduced the function of *Opa1*-mut and *TFAM*-OE clock neurons (Fig. 3B, D), suggesting that increasing mitochondrial NAD^+^ regeneration not only does not restore function but may rather further perturb the metabolic state of these neurons. (See Discussion.)

In addition, we used *Ndi1*, a non-proton-pumping yeast NADH dehydrogenase that oxidizes NADH to NAD^+^ and transfers electrons to ubiquinone (Bakker et al., 2001). *Ndi1* expression failed to replace LDH function in *Opa1*-mut or *TFAM*-OE neurons (Fig. S2A, B), further supporting the above indications that NAD^+^ levels alone are not key (Fig. 3). To make sure that *Ndi1* is functional in our system, we reproduced a previous study showing that early lethality caused by pan-neuronal loss of mitochondrial Complex I function using *nSyb-Gal4* is rescued by *Ndi1* expression (Fig. S2C, Granat et al., 2023). However, *Ndi1* expression did not rescue developmental lethality caused by other pan-neuronal mitochondrial issues including *Opa1-*mut, RNAi-mediated *Opa1* knockdown, or *TFAM-OE* (Fig. S2C), confirming that *Ndi1* is not sufficient to rescue several other broad forms of mitochondrial dysfunction.

### mitoLbNOX expression alone causes mitochondrial dysfunction and a transcriptional response

Because expression of both *LbNOX* and *mitoLbNOX* modified neuronal function in mitochondrial dysfunction backgrounds (Fig. 3B, D), we examined the expression and localization of LbNOX via its FLAG tag. As predicted, LbNOX showed diffuse FLAG staining throughout the cytoplasm and nucleus and did not alter mitochondrial morphology as visualized by mitoGFP (Fig. 4A, S3). In contrast, mitoLbNOX-FLAG not only localized to mitochondria but also caused their fragmentation (Fig. 4A, S3).

**Figure 4:**
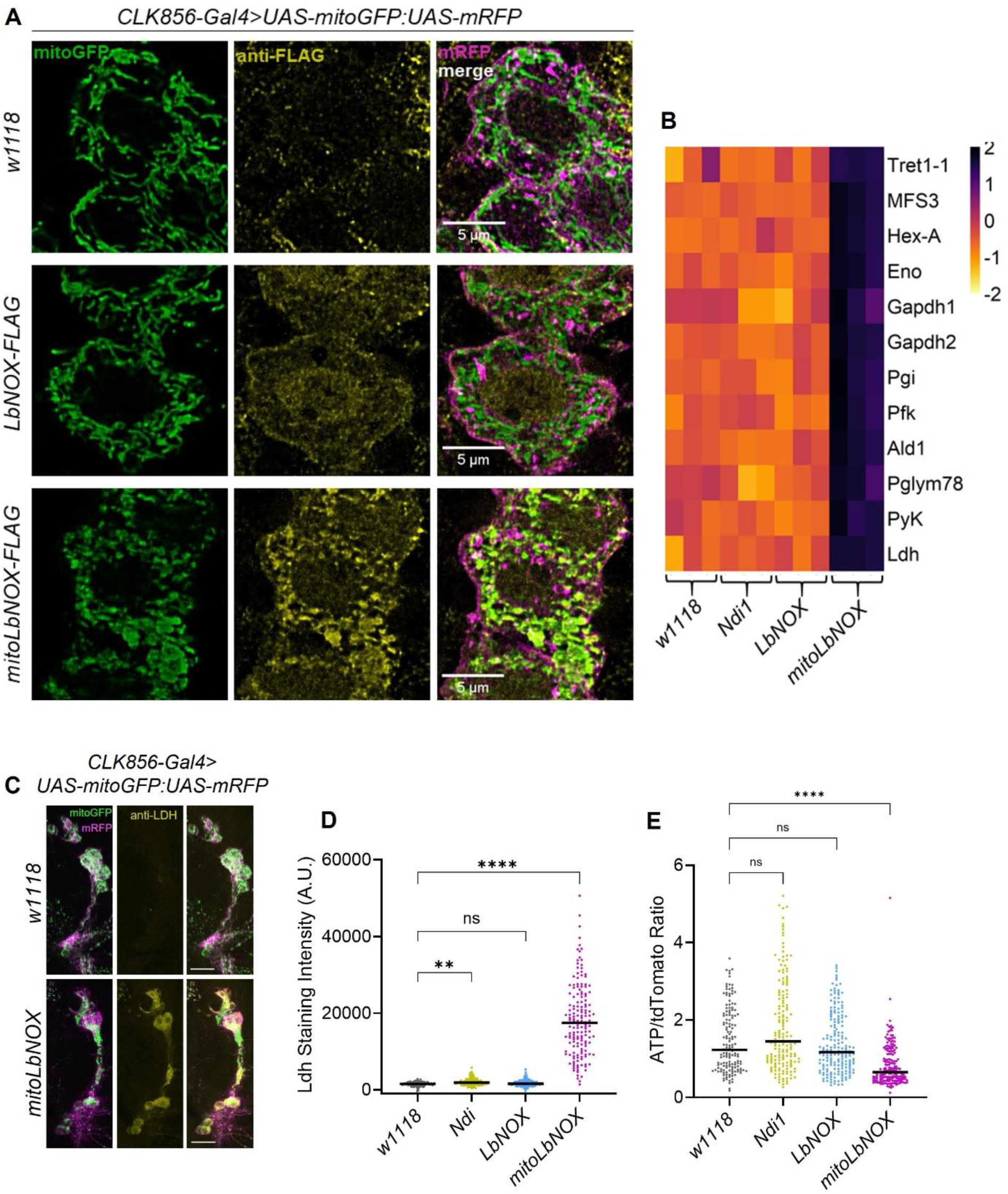
mitoLbNOX, but not LbNOX, expression leads to mitochondrial fragmentation and upregulation of glycolytic genes. **A.** Representative images of large-PDF neurons, a subset of clock neurons marked by *CLK856-Gal4*, from ∼15-day-old flies with mitochondria labeled using *UAS-mitoGFP* and *UAS-mRFP*. Anti-FLAG staining is dispersed throughout the cell and around the nucleus in LbNOX-expressing neurons, whereas it localizes to mitochondria in mitoLbNOX-expressing neurons. Mitochondrial morphology is also fragmented in mitoLbNOX-expressing neurons. **B.** Heatmap showing expression levels of genes involved in glycolysis and sugar transport across three replicates per genotype. Glycolytic genes and sugar transporters are upregulated specifically in mitoLbNOX-expressing clock neurons. **C.** Representative images of LDH staining in ventral clock neurons from ∼15-day-old flies of the indicated genotypes. Clock neurons were visualized using *UAS-mitoGFP* and *UAS-mRFP*. Scale bars represent 20 μm. **D.** Quantification of LDH staining intensity; n ≥ 117 neurons from ≥10 hemibrains per genotype. Adjusted P > 0.05 is denoted as not significant (ns); **P < 0.01, ****P < 0.0001, Kruskal–Wallis test followed by Dunn’s post hoc test. **E.** Ratio of *iATPSnFR* to *tdTomato* in ∼15-day-old lateral clock neurons (sLNvs, lLNvs, and LNds) of the indicated genotypes; n ≥ 150 cell regions from ≥10 brains per genotype. Expression of mitoLbNOX results in a significant reduction in ATP levels. Kruskal–Wallis test followed by Dunn’s post hoc test; ****P < 0.0001; ns, not significant (P > 0.05).

We next examined the transcriptomic response of clock neurons to these NADH-oxidizing enzymes at 5 weeks of adulthood, the same age used for the transcriptomic analysis of *Opa1*-mut (Richhariya et al., 2025) and *TFAM*-OE neurons (Fig. 2). Expression of *Ndi1* or *LbNOX* did not substantially alter the transcriptome as there were very few differentially regulated genes (Fig. S4A-B), but expression of *mitoLbNOX* induced a large transcriptomic response which is reminiscent of the cancer-like transcriptomic profile of *Opa1*-mut clock neurons (Fig. S4C, Richhariya et al., 2025), including a much greater number of upregulated than downregulated genes (Fig. S4C). Several glycolytic genes including *Ldh* were upregulated in *mitoLbNOX-*expressing neurons (Fig. 4B, S4D). Not unexpectedly, *mitoLbNOX*-expressing clock neurons had a concurrent increase in LDH protein levels (Fig. 4C, D). These neurons also had lower ATP levels (Fig. 4E), indicating compromised mitochondrial function. This is similar to the loss of fusion (Richhariya et al., 2025) and consistent with their mitochondrial fragmentation (Fig. 4A, S3).

Despite these similarities, there was an important distinction in the transcriptomic response to the different mitochondrial perturbations. *ND-75* knockdown and mitoLbNOX expression both caused increased levels of mitochondrial DNA-encoded transcripts (Fig. S5A, B), perhaps indicating a compensatory response to weak mitochondrial activity. This is different from *Opa1*-mut clock neurons, which show reduced expression of mitochondrial transcripts (Richhariya et al., 2025), likely reflecting in this case a reduction in mtDNA copy number in the absence of fusion. Interestingly, *TFAM*-OE neurons are different yet again: some mtDNA transcripts were upregulated whereas others were downregulated (Fig. S5C), reflecting an imbalance in mtDNA transcription. Because glycolytic gene upregulation occurs despite these distinct effects on mitochondrial transcript expression, the glycolytic response is probably not triggered by a mtDNA-encoded gene expression change. On the other hand, all these perturbations upregulate the transcription factor ATF4. Although this occurs to different extents (Fig. S5D-F), these data together with our previous experiments (Richhariya et al., 2025) suggest that induction of ATF4 contributes to the glycolytic gene upregulation and represents a shared response to mitochondrial dysfunction. (See Discussion).

### LDH protects mitoLbNOX-expressing neurons from neurodegeneration

To address the functional consequence of *Ldh* upregulation in *mitoLbNOX*-expressing neurons, we examined flies in which clock neurons expressed *mitoLbNOX* and were also *Ldh*-mut. As expected, this double perturbation caused severely reduced rhythmicity (Fig. 5A). This was much more severe than *mitoLbNOX* expression alone (Fig. 3A) and indicates that LDH compensates for the loss of neuronal function in *mitoLbNOX-*expressing neurons. Note that this is specific to *mitoLbNOX* as there is no comparable interaction between *LbNOX* and *Ldh*, as might be expected from the lack of a strong transcriptional response to *LbNOX* expression (Fig. S4B, D). Although there was no visible neurodegeneration in 40-day-old flies due to *mitoLbNOX* expression, the addition of *Ldh*-mut to this genotype caused neurodegeneration (Fig. 5B, C). The results indicate that LDH upregulation and likely glycolysis more generally protect neurons from mitoLbNOX-induced dysfunction and neurodegeneration.

**Figure 5:**
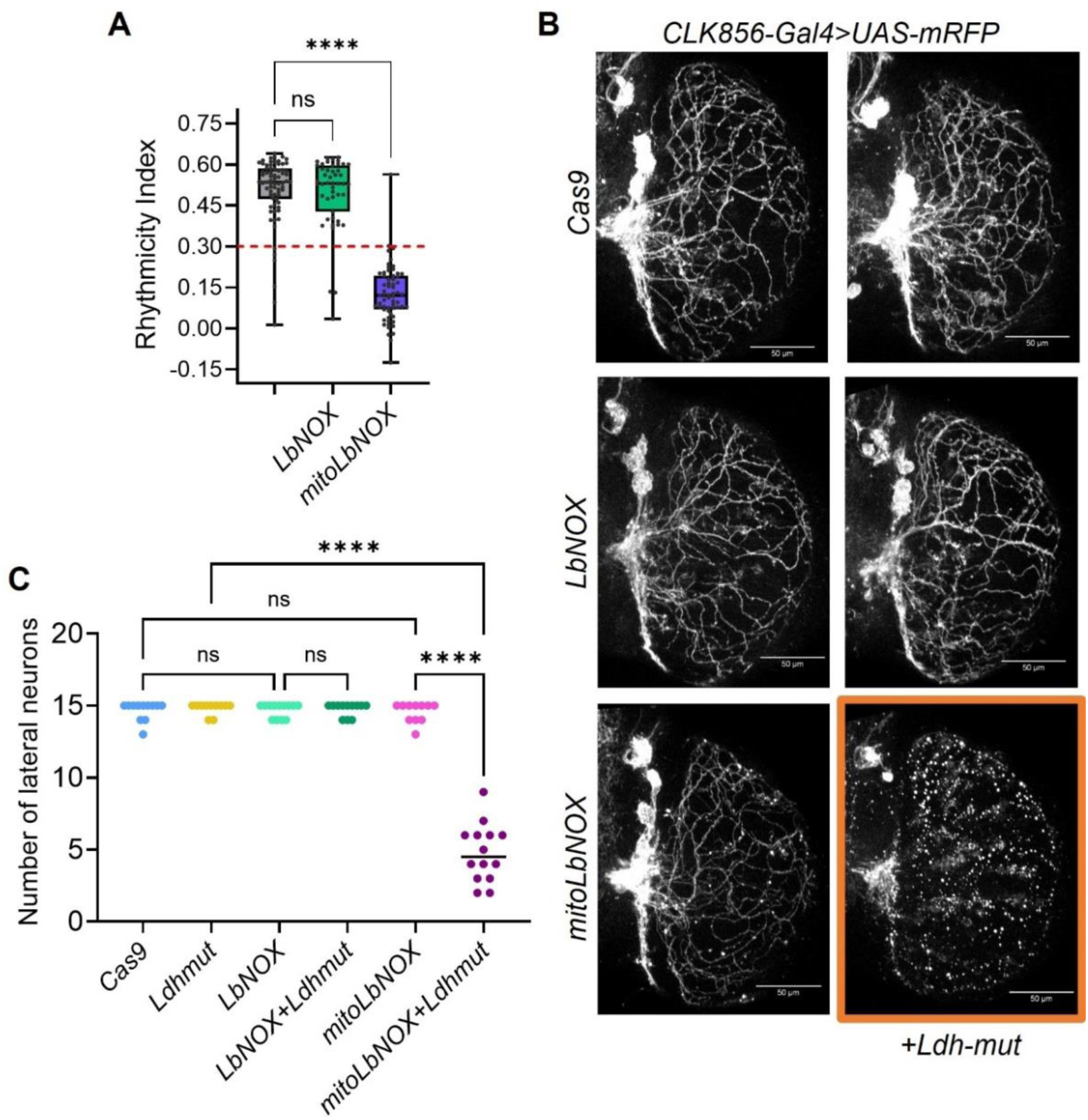
*Ldh* preserves neuronal function and prevents neurodegeneration in mitoLbNOX-expressing clock neurons. **A.** Rhythmicity index (RI), a measure of clock neuron function, for *Ldh-mut* flies expressing the indicated transgenes in clock neurons using *CLK856-Gal4* from ∼3wk old flies. Expression of LbNOX had no effect, whereas expression of mitoLbNOX in combination with *Ldh-mut* led to a severe loss of rhythmicity. RI < 0.3, indicated by the red dotted line, is considered arrhythmic. ****P < 0.0001; ns, not significant (P > 0.05), Kruskal–Wallis test followed by Dunn’s post hoc test. **B.** Representative images of ventral clock neurons and their projections in ∼40-day-old brains, labeled with *UAS-mRFP* driven by *CLK856-Gal4*. Neurons express the transgenes indicated on the left; those in the right column are additionally *Ldh-mut*. Neurodegeneration is evident in mitoLbNOX + *Ldh-mut* clock neurons. Scale bars represent 50 μm. **C.** Quantification of the number of ventral clock neurons (sLNvs, lLNvs, and LNds) per hemibrain for the genotypes shown in (B). Only mitoLbNOX + *Ldh-mut* flies had significantly fewer neurons than either single perturbation. n ≥ 11 hemibrains per genotype. ****P < 0.0001; ns, not significant (P > 0.05), Kruskal–Wallis test followed by Dunn’s post hoc test.

## Discussion

Neurons are post-mitotic, long-lived and morphologically unusual. They therefore experience and must cope with metabolic challenges in special ways. Our previous work described the response to the absence of mitochondrial fusion in *Opa1*-deficient neurons (Richhariya et al., 2025), and we show here that glycolytic upregulation is a general response to diverse sources of mitochondrial dysfunction. They include *TFAM* overexpression, *mitoLbNOX* expression and reduced Complex I function. For Complex I knockdown, glycolytic upregulation occurs despite indications that substantial mitochondrial function remains. The transcriptional response includes *Ldh* upregulation, and LDH is required to maintain function in *TFAM-*OE and *mitoLbNOX*-expressing clock neurons. LDH is also necessary to prevent the neurodegeneration that is a consequence of mitochondrial dysfunction. Importantly, our experiments indicate that this LDH-mediated neuroprotection is due to increased glycolytic activity rather than simply increasing NAD^+^/NADH levels.

### Previous indications of a glycolytic response due to mitochondrial dysfunction

Neuronal *TFAM* overexpression in *Drosophila* increases *Ldh* expression (Cagin et al., 2015). Similarly, Complex I knockdown in broad sets of *Drosophila* neurons causes upregulation of *Ldh* (Granat et al., 2023), and neuronal Complex IV deficiency in mice increases glycolytic metabolism (Garcia et al., 2022). In non-neuronal cells, Complex IV disruption in *Drosophila* cells induces glycolytic gene expression (Freije et al., 2012; Sorge et al., 2020), whereas Complex I dysfunction drives a similar glycolytic switch in mammalian smooth-muscle cells (Rafikov et al., 2015). These studies did not test whether the glycolytic response protects cells from mitochondrial stress.

There was however some indication of function from a study in *Drosophila* eye discs, in which *Ldh* knockdown exacerbated the developmental delay caused by Complex I knockdown (Veits et al., 2026). Moreover, glial cells with impaired Complex IV function can survive long term with upregulated glycolysis (Supplie et al., 2017), and Pink1-deficient MEFs can sustain proliferation through enhanced glycolysis (Requejo-Aguilar et al., 2014). Nonetheless, the functional importance of glycolytic upregulation in response to ETC and mitochondrial dysfunction in flies and mammals, particularly in post-mitotic neurons, had not been well-substantiated.

How does mitochondrial dysfunction signal to increase cytoplasmic glycolysis? The integrated stress response transcription factor ATF4 has been shown to be upregulated in *TFAM*-OE neurons (Hunt et al., 2019). This is similar to our results in mitoLbNOX-expressing neurons (Fig. S5D). ATF4 is also upregulated and important for the glycolytic upregulation in *Opa1*-mut neurons (Richhariya et al., 2025). In myotubes, ATPase inhibition activates the integrated stress response (ISR) through a mechanism related to mitochondrial inner-membrane hyperpolarization (Mick et al., 2020). This raises the possibility that a similar mitochondrial stress mechanism contributes to ATF4 induction and compensatory glycolysis in clock neurons experiencing mitochondrial dysfunction.

### Elevating neuronal NAD⁺/NADH ratio cannot compensate for mitochondrial stress

LDH catalyzes the conversion of pyruvate to lactate while regenerating NAD⁺ from NADH. Might the role of LDH then be to maintain sufficient NAD in the face of mitochondrial dysfunction? To test this hypothesis, we used multiple genetic strategies to increase this ratio.

They included expression of the yeast NADH dehydrogenase *Ndi1*. It can rescue the shortened lifespan of flies with neuronal Complex I deficiency (Granat et al., 2023). Consistent with this result, *Ndi1* also rescued the developmental lethality caused by Complex I knockdown (Fig. S2C). Thus, *Ndi1* can functionally replace Complex I when downstream respiratory chain components are functional. However, *Ndi1* did not rescue the developmental lethality of other mitochondrial perturbations, i.e., *Opa1* knockdown, knockout or TFAM overexpression (Fig. S2C). It also failed to replace LDH function in *Opa1*-deficient or *TFAM-OE* clock neurons (Fig. S2A-B). Although restoration of mitochondrial NADH oxidation and redox homeostasis was proposed to contribute to the rescue of some phenotypes caused by Complex I deficiency (McElroy et al., 2020), *Ndi1* can also support electron flow through the respiratory chain and contribute to oxidative phosphorylation (Jiménez-Gómez et al., 2023). The latter may be a better explanation given its failure to rescue broader forms of mitochondrial dysfunction.

Expression of *LbNOX* or mitochondria-targeted *mitoLbNOX* was another strategy we used to oxidize NADH and increase the NAD⁺/NADH ratio in *Opa1-mut* or *TFAM-OE* backgrounds. Surprisingly, both slightly but significantly reduced the rhythmicity of these already compromised flies (Fig. 3B, D). A plausible explanation is that a decrease in NADH would decrease glycolysis by limiting LDH-dependent pyruvate-to-lactate conversion; this reaction requires NADH as a cofactor. MitoLbNOX can also lower cytosolic NADH levels (Titov et al., 2016), likely through redox shuttles, and could therefore similarly limit LDH-dependent compensation in mitochondria-deficient neurons. However, LbNOX and mitoLbNOX both failed to replace LDH in mitochondria-defective clock neurons (Fig. 3C, E), so an increase in NAD⁺/NADH levels alone may be insufficient to account for the potent LDH-mediated rescue.

These results contrast with the rescue of mitochondrial defects by LbNOX in cultured cells (Diebold et al., 2019; Titov et al., 2016) and the partial rescue of Complex I loss in proliferating eye disc cells (Veits et al., 2026). In neurons, the interaction between mitochondrial defects and NADH oxidation is less consistent. In mice heterozygous for the *Opa1^V291D^*mutation, retinal ganglion cells had lower glycolysis and mitoLbNOX expression improved function and survival (Kang et al., 2026). However, mitoLbNOX failed to rescue cellular or behavioral defects caused by Complex I dysfunction in dopaminergic neurons (D’Alessandro et al., 2025). These differences may in part reflect the different roles of glycolysis in dividing and post-mitotic cells.

### mitoLbNOX as a mitochondrial stressor

A surprising finding was the mitochondrial defect caused by the targeting of *LbNOX* expression to the mitochondria in otherwise wild-type neurons. NADH is the primary electron donor to the electron transport chain (ETC), and its oxidation to NAD^+^ through an ETC-independent pathway might well reduce electron flux through the ETC. Consistent with this possibility, ATP levels were decreased in mitoLbNOX-expressing neurons (Fig. 4E). The accompanying compensatory increase in glycolysis may reflect activation of an ATF4-mediated ISR secondary to mitochondrial dysfunction (Mick et al., 2020; Richhariya et al., 2025). This is similar to the glycolytic response following reduced complex I activity (Fig. S1) and TFAM overexpression (Fig. 2).

Importantly, the comparable expression of untargeted LbNOX had little effect on mitochondrial or neuronal function; the protein was distributed throughout the cytosol and nucleus, ATP levels remained normal and the transcriptome was largely unchanged (Fig. 4, S4B). Therefore, increasing the NAD^+^/NADH ratio within neuronal mitochondria is substantially more detrimental than raising the ratio in the cytoplasm. This compartment-specific sensitivity is consistent with the much lower NAD^+^/NADH ratio normally maintained in mitochondria relative to the cytoplasm (Cambronne and Kraus, 2020).

We note that short-term expression of mitoLbNOX in HeLa cells for 24 hours did not alter mitochondrial morphology (Titov et al., 2016), suggesting that prolonged expression, higher expression levels, or the *in vivo* neuronal context may be required for the observed adverse effects on mitochondrial health. In any case, these findings suggest caution when using mitoLbNOX as a positive intervention, especially for prolonged expression in neurons.

## Materials and Methods

### Fly rearing and stocks

Flies were raised on standard cornmeal medium supplemented with yeast at 25°C under 12h light:12h dark conditions. For experiments using aged-flies, ∼0-4 days-old male flies were collected and separated into vials containing 30-40 flies. Aging flies were transferred to fresh food every 3-4 days until ready for experiments. For knockdown or knockout of glycolysis genes, preference was given to RNAi lines generated using the Valium 20 vector which has strong somatic expression (Perkins et al., 2015). If not available, gRNA lines were used. These gRNA lines have only one gRNA targeting the gene and are expressed ubiquitously; specificity is achieved by UAS-Cas9. All fly lines used are listed in Table S1.

### Generation of fly lines

To generate the *UAS-LbNOX* and *UAS-mitoLbNOX*, the coding sequences along with FLAG tags (Titov et al., 2016) were synthesized and inserted into the 20X-UAS vector pJFRC7 cut with XhoI and XbaI by Genscript (Piscataway, NJ, USA). The plasmid sequences were verified by whole plasmid sequencing performed by Plasmidsaurus using Oxford Nanopore Technology. Fly lines were generated by injecting plasmids into embryos (BDSC 9744) with insertion on the third chromosome by BestGene Inc. (Chino Hills, CA, USA).

### Circadian Behavior Assay

Circadian behavior was assayed as described previously (Richhariya et al., 2023). Briefly, male flies, at ∼3 weeks were loaded into Drosophila Activity Monitor (DAM) tubes containing sucrose food (4% sucrose and 2% agar). Because *Opa1+Ldh-mut* had a strong phenotype at 3 weeks (Richhariya et al., 2025), the same age was used to assay additional interactions.

Light boxes with programmable LED light intensities were used to control light conditions and were kept inside a temperature-controlled incubator set to 25°C. Post entrainment to a 12 hours light: 12 hours dark cycle for at least 3 days, flies were switched to constant darkness for at least 7 days. Rhythmicity index, used as a measure of the strength of the circadian rhythm was calculated for constant darkness (DD) days 2-7 using Sleep and Circadian Analysis MATLAB Program (SCAMP) developed by Christopher G. Vecsey (Vecsey et al., 2024). Percent rhythmic flies reported in Table 1 are the percentage of flies with RI values > 0.3. Period values were analyzed and reported only from rhythmic flies (RI>0.3) were used. Rhythmicity Index and Period data are presented as box plots showing all data points, and whiskers extending from min to max.

### Longevity assay

Male flies were collected at 0-3d old, ∼25 flies were placed in a vial and transferred to fresh food every 3-4 days and dead flies counted. At least 90 flies were tested per genotype.

### Immunohistochemistry

Immunostaining for *Drosophila* brains was performed as described previously (Richhariya et al., 2023). Briefly, flies were fixed in 4% PFA (Fisher Scientific #50-980-487) and then brains were dissected and blocked in blocking buffer (10% Normal Goat Serum (NGS, Jackson Labs #005-000-121) in 0.5% PBST). The following primary antibodies in blocking buffer were used: chicken anti-GFP (1:2000, Abcam #ab13970), rat anti-RFP (1:1000, Proteintech #5f8), rabbit anti-LDH (1:500, Boster Bio #DZ41222), rabbit anti-FLAG (1:250, Proteintech 80801-2-RR). Primary antibody incubations were done for 15-18h at 4°C for GFP and RFP or ∼48h at 4ᴼC if LDH or FLAG was used. Secondary antibodies were used at 1:500 dilution in blocking buffer and were incubated either for 3 hours at RT or 4°C overnight. All samples compared to each other were processed together. Brains were mounted in Vectashield PLUS (Vector Laboratories #H-1900).

### Image acquisition and analysis

All images were acquired on Leica Stellaris 8 confocal microscope equipped with a white light laser as described in (Richhariya et al., 2025). Imaging settings were constant across different samples in a set. Image processing and analysis were performed using Fiji (Schindelin et al., 2012).

For visualizing mitochondrial morphology and FLAG-localization, images were acquired using a 63X oil objective with NA of 1.4 with additional optical zoom. Deconvolution was performed on acquired images using LIGHTNING on the LAS X software.

For visualizing the projections of ventral clock neurons, images were acquired of the optic lobe region with the 20X air objective with an NA of 0.75 with additional optical zoom. Images were adjusted for brightness/contrast, with the same level of correction applied to all the samples in an experiment. LDH levels were quantified as described in (Richhariya et al., 2025).

ATP measurements were done and quantified as described in (Richhariya et al., 2025). For lactate measurements, brains expressing the *CanlonicSF* sensor and *tdTomato* under *CLK856-Gal4* were dissected from ∼3-week-old male flies in adult hemolymph-like (AHL) saline supplemented with 5 mM glucose. Brains were maintained at room temperature in AHL + glucose and, within 15 min of dissection, mounted anterior side up on a slide in AHL with the coverslip sealed. Brains were imaged immediately using a 20X-air objective with sequential scan settings. Quantification was performed on ROIs using the Time Series Analyzer v3 plugin in Fiji, and values are reported per cell-body region. All data points are plotted with median as a line.

### FACS-sorting and generation of sequencing libraries

Two sequencing datasets were generated as part of this study, both from ∼35-day-old male flies at ZT14. The first set consisted of control (*CLK856-Gal4>w1118*), Complex I knockdown (*CLK856-Gal4>UAS-ND-75 RNAi)* and TFAM-overexpression (*CLK856-Gal4>UAS-TFAM*) in triplicates. The same control set was used for complex I KD and *TFAM*-OE and these three genotypes were processed and sequenced together. The second set consisted of control (*CLK856-Gal4>w1118*), and expression of NAD^+^/NADH modulating enzymes - Ndi1 (*CLK856-Gal4>UAS-Ndi1*), LbNOX (*CLK856-Gal4>UAS-LbNOX) and* mitoLbNOX *(CLK856-Gal4>UAS-mitoLbNOX)* in triplicates. In addition to these transgenes, each genotype also contained *UAS-mitoGFP* and *UAS-RFP.* Clock neurons were FACS sorted using the GFP channels as described in (Richhariya et al., 2025). From sorted neurons, smart-seq3 based sequencing libraries were generated using protocols described in (Richhariya et al., 2025). Libraries were sequenced on Illumina Nextseq1000 with paired end read length of 75 X 45.

### Sequencing data analysis

The reads were mapped to the *Drosophila* genome (dm6) using the zUMIs pipeline (Parekh et al., 2018). UMI counts mapping to exons of each gene were extracted before further analysis. Data were normalized using DESeq2 (Love et al., 2014). Volcano plots were generated from the DESeq2 output using the package EnhancedVolcano (Blighe K, Rana S, Lewis M, 2023). DESeq2 output normalized counts were transformed using normTransform and then used to create heatmaps of selected genes with pheatmap (Kolde R, 2019). Heatmaps were scaled for rows and not clustered for rows or columns.

## Acknowledgements

We thank all members of the Rosbash lab for helpful discussions. We thank Dr. Thomas R. Clandinin for sharing the *UAS-iATPSnFR* fly line and Dr. Joseph Bateman for sharing the *UAS-TFAM* line. Fly stocks obtained from the Bloomington Drosophila Stock Center (NIH P40OD018537) were used in this study. This work was supported by the Howard Hughes Medical Institute (HHMI).

---

**Figure S1:**
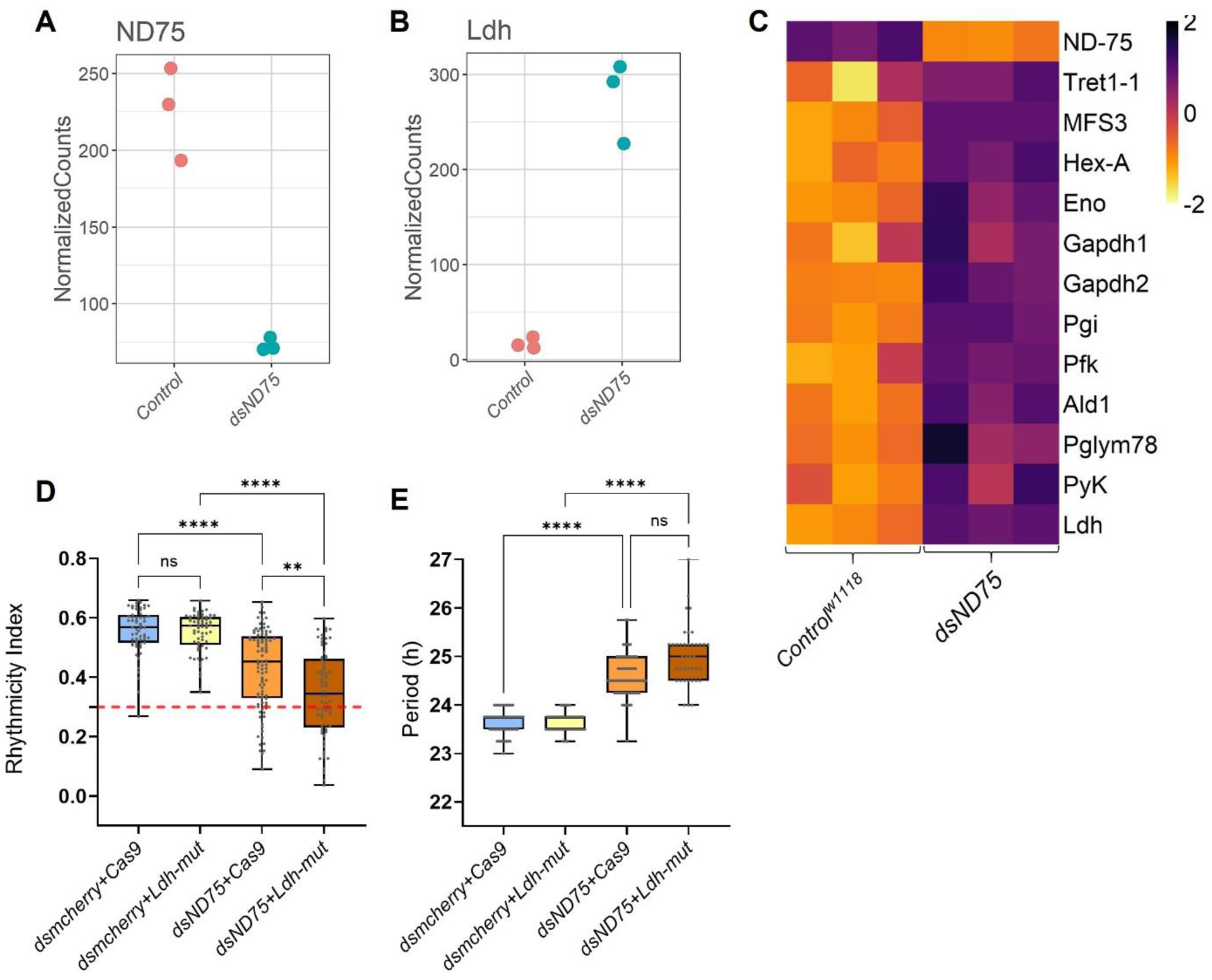
The glycolysis pathway is upregulated in Complex I knockdown clock neurons. **A–B.** Normalized counts for *ND-75* (A) and *Ldh* (B) from bulk sequencing of ∼35-day-old control and *ND-75* knockdown (*dsND-75*) clock neurons, confirming *ND-75* knockdown (A) and showing strong *Ldh* upregulation (B). **C.** Heatmap showing expression levels of genes involved in glycolysis and sugar transport across three replicates per genotype. Glycolytic genes and sugar transporters show a general increase in expression upon *ND-75* knockdown; *ND-75* levels are shown in the top row. **D–E.** Rhythmicity index (RI), a measure of clock neuron function (D), and circadian period (E) for the indicated genotypes from ∼3-4-week-old flies. RI < 0.3, indicated by the red dotted line, is considered arrhythmic. *ND-75* knockdown in clock neurons causes a modest but significant reduction in rhythmicity and a longer circadian period. Combined *dsN-D75* and *Ldh-mut* perturbation results in significantly lower rhythmicity than either single perturbation, although the overall effect is smaller than that observed for *TFAM-OE + Ldh-mut* (Fig. 2E). n ≥ 58 flies per genotype. **P < 0.01; ****P < 0.0001; ns, not significant (P > 0.05), Kruskal–Wallis test followed by Dunn’s post hoc test.

**Figure S2:**
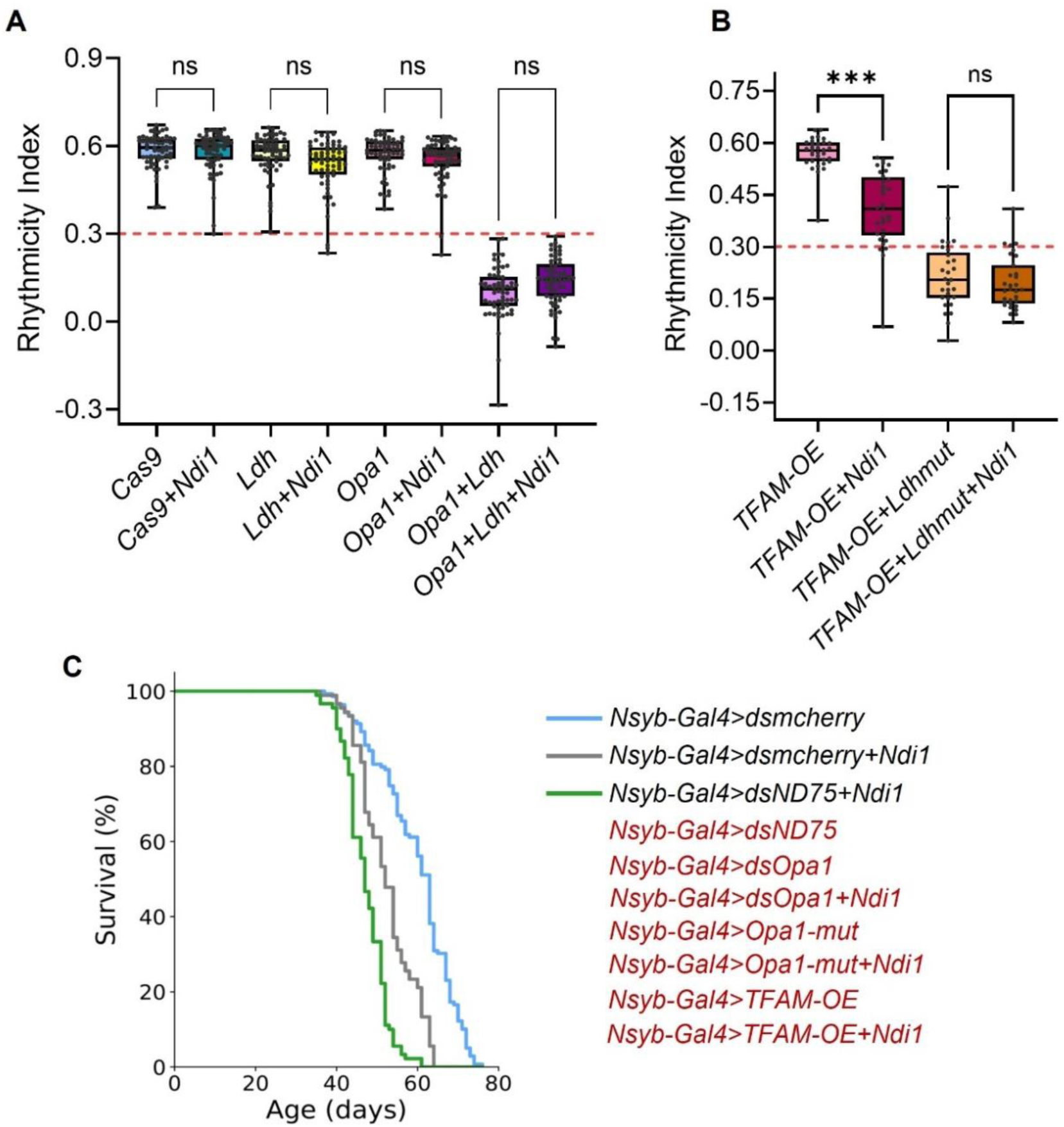
Ndi1 rescues Complex I deficiency but not general mitochondrial dysfunction. **A–B.** Rhythmicity index (RI), a measure of clock neuron function, for the indicated genotypes driven by *CLK856-Gal4* from ∼3-week-old flies. RI < 0.3, indicated by the red dotted line, is considered arrhythmic. n ≥ 29 flies per genotype. ***P < 0.001; ns, not significant (P > 0.05), Kruskal–Wallis test followed by Dunn’s post hoc test. *Ndi1* does not rescue the rhythmicity defects of *Opa1 + Ldh-mut* or *TFAM-OE + Ldh-mut*. *Ndi1* causes a modest but significant reduction in rhythmicity in *TFAM-OE*-expressing clock neurons. **C.** Survival curves of male flies of the indicated genotypes. No adult flies were obtained for genotypes indicated in red, consistent with developmental lethality. *Ndi1* rescues developmental lethality and restores substantial lifespan in flies with pan-neuronal Complex I knockdown using *Nsyb-Gal4* (*dsND-75*), but not in flies with *Opa1-KD*, *Opa1-mut*, or *TFAM-OE.* n ≥ 90 flies per genotype.

**Figure S3:**
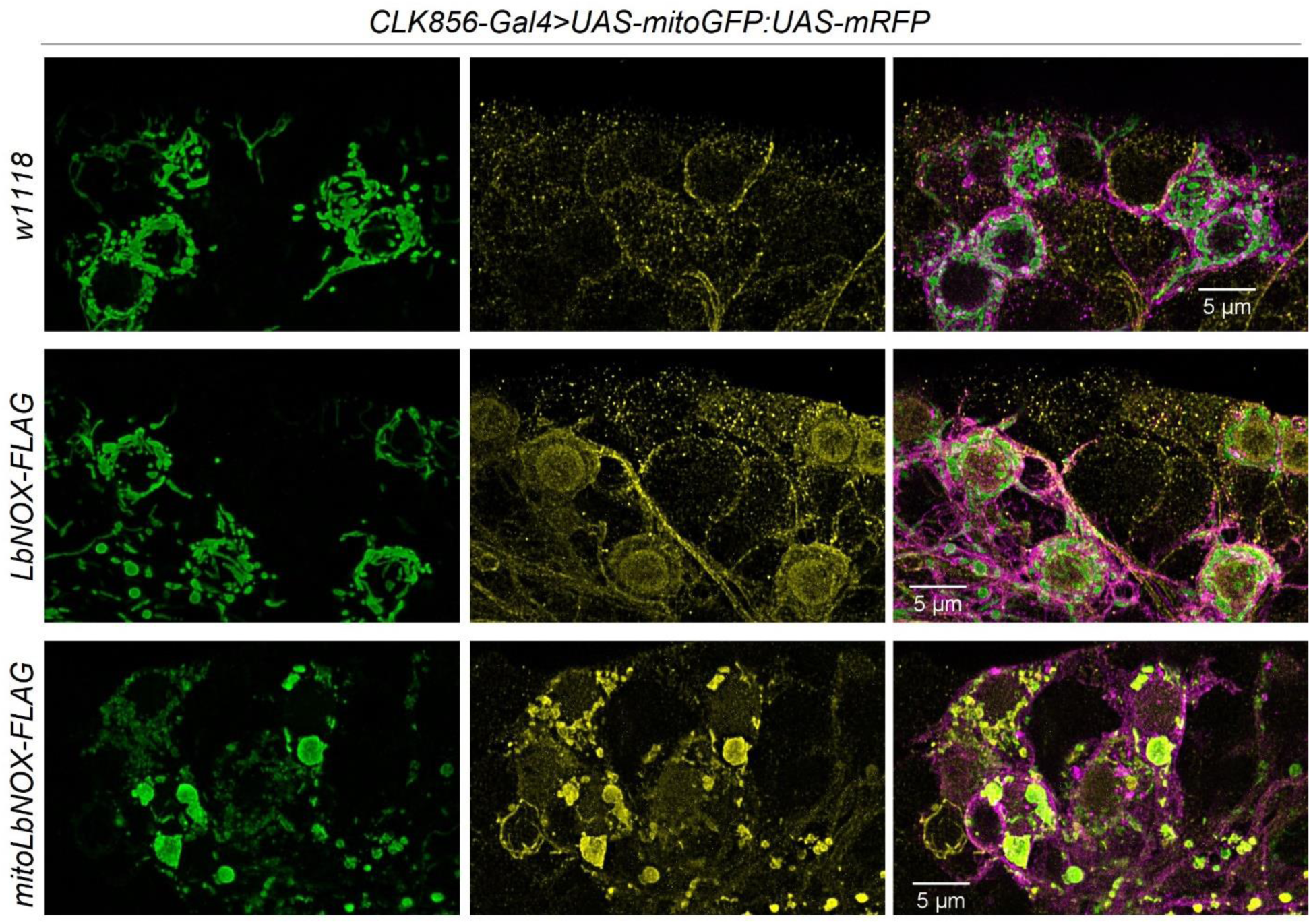
mitoLbNOX localizes to the mitochondria and causes mitochondrial fragmentation in dorsal DN1 neurons. Representative images of DN1 neurons, a subset of clock neurons from ∼15-day-old flies marked by *CLK856-Gal4*, with mitochondria labeled using *UAS-mitoGFP* and *UAS-mRFP*. Anti-FLAG staining is dispersed throughout the cell and around the nucleus in LbNOX-expressing neurons, whereas it localizes to mitochondria in mitoLbNOX-expressing neurons. Mitochondrial morphology is also fragmented in mitoLbNOX-expressing neurons.

**Figure S4:**
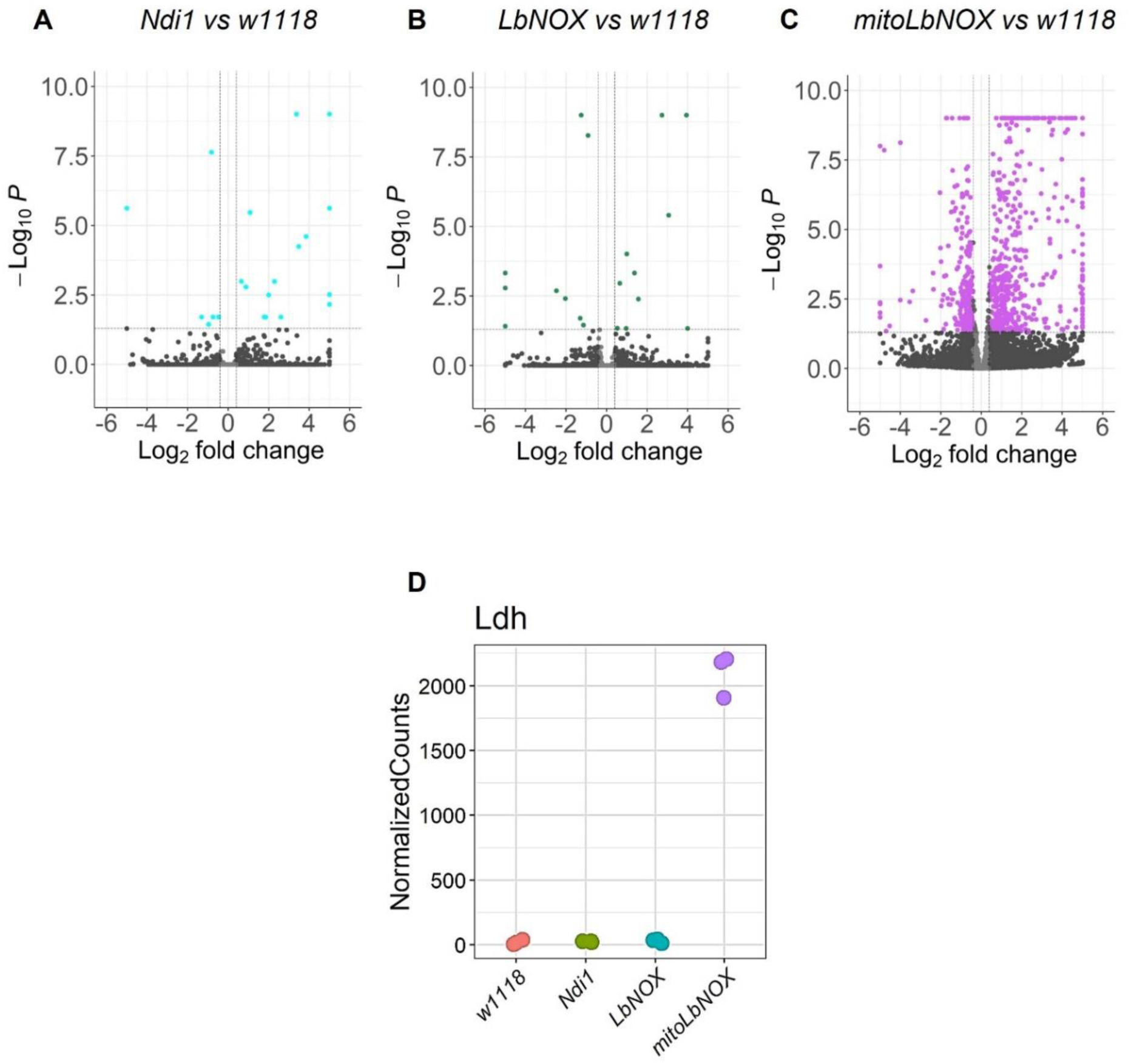
*mitoLbNOX* expression elicits a strong transcriptomic response, whereas *Ndi1* or *LbNOX* expression does not. **A–C.** Volcano plots of ∼35-day-old clock neurons expressing NAD^+^/NADH-manipulating enzymes compared with *w1118* controls. Colored dots indicate significantly differentially expressed genes (adjusted *P* < 0.05). Only *mitoLbNOX* expression triggers a strong transcriptomic response, with more genes upregulated than downregulated. **D.** Normalized counts for *Ldh* from the same bulk sequencing dataset show *Ldh* upregulation only in *mitoLbNOX-*expressing clock neurons.

**Figure S5:**
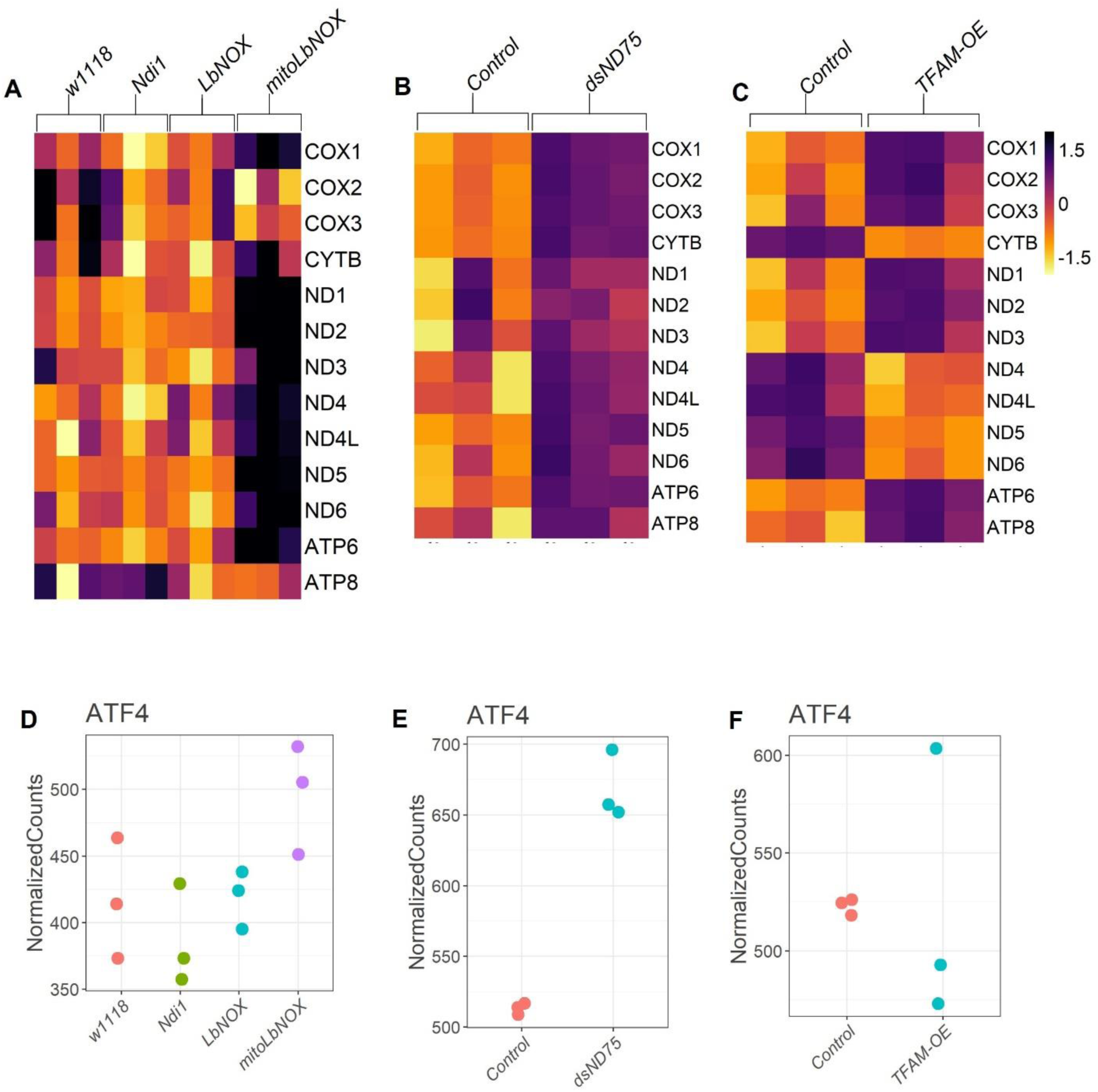
Regulation of mitochondrial transcripts differs across mitochondrial perturbations, whereas ATF4 is upregulated across several mitochondrial perturbations. **A–C.** Heatmaps showing expression levels of genes encoded by the mitochondrial genome in the indicated genotypes. Whereas mitoLbNOX expression and Complex I knockdown using *dsND-75* cause a general upregulation of mitochondrial genes, *TFAM-OE* causes upregulation of some mitochondrial genes and downregulation of others. **D-F.** Normalized counts for *ATF4* from clock neurons of indicated genotypes.

**Table S1:**
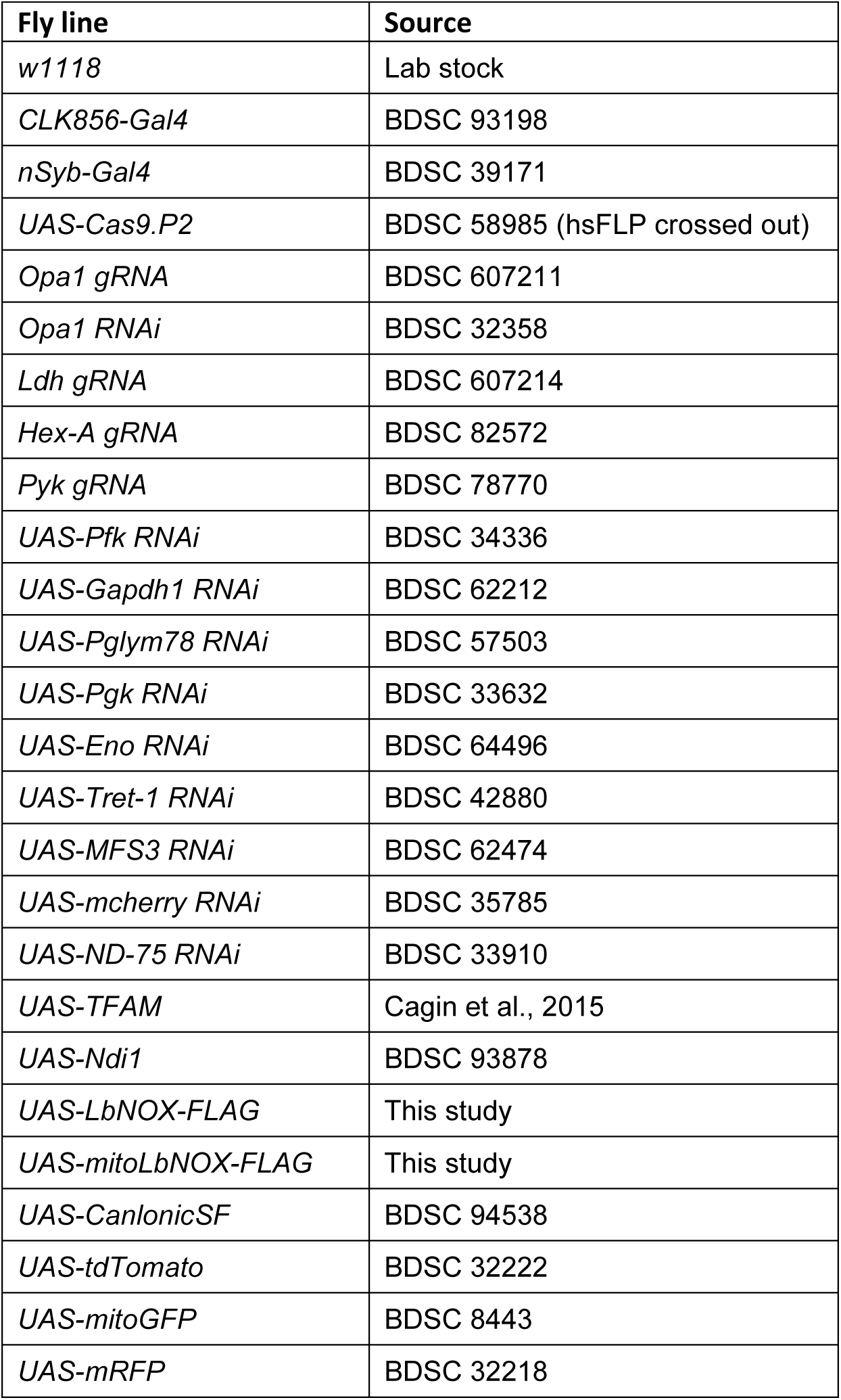
List of fly lines used.

## References

Aburto C, Galaz A, Bernier A, Sandoval PY, Holtheuer-Gallardo S, Ruminot I, Soto-Ojeda I, Hertenstein H, Schweizer JA, Schirmeier S, Pástor TP, Mardones GA, Barros LF, San Martín A. 2022. Single-Fluorophore Indicator to Explore Cellular and Sub-cellular Lactate Dynamics. ACS Sensors 7:3278–3286. DOI: 10.1021/acssensors.2c00731

Bahadorani S, Cho J, Lo T, Contreras H, Lawal HO, Krantz DE, Bradley TJ, Walker DW. 2010. Neuronal expression of a single-subunit yeast NADH-ubiquinone oxidoreductase (Ndi1) extends Drosophila lifespan. Aging Cell 9:191–202. DOI: 10.1111/j.1474-9726.2010.00546.x, PMID: 20089120

Bakker BM, Overkamp KM, van Maris AJ null, Kötter P, Luttik MA, van Dijken JP null, Pronk JT. 2001. Stoichiometry and compartmentation of NADH metabolism in Saccharomyces cerevisiae. FEMS microbiology reviews 25:15–37. DOI: 10.1111/j.1574-6976.2001.tb00570.x, PMID: 11152939

Betarbet R, Sherer TB, MacKenzie G, Garcia-Osuna M, Panov AV, Greenamyre JT. 2000. Chronic systemic pesticide exposure reproduces features of Parkinson’s disease. Nature Neuroscience 3:1301–1306. DOI: 10.1038/81834

Blighe K, Rana S, Lewis M. 2023. EnhancedVolcano: Publication-ready volcano plots with enhanced colouring and labeling_. R package version 1.13.2,.

Bonekamp NA, Jiang M, Motori E, Garcia Villegas R, Koolmeister C, Atanassov I, Mesaros A, Park CB, Larsson N-G. 2021. High levels of TFAM repress mammalian mitochondrial DNA transcription in vivo. Life Science Alliance 4:e202101034. DOI: 10.26508/lsa.202101034, PMID: 34462320

Cagin U, Duncan OF, Gatt AP, Dionne MS, Sweeney ST, Bateman JM. 2015. Mitochondrial retrograde signaling regulates neuronal function. Proceedings of the National Academy of Sciences of the United States of America 112:E6000–E6009. DOI: 10.1073/pnas.1505036112, PMID: 26489648

Camandola S, Mattson MP. 2017. Brain metabolism in health, aging, and neurodegeneration. The EMBO Journal 36:1474–1492. DOI: 10.15252/embj.201695810, PMID: 28438892

Cambronne XA, Kraus WL. 2020. Location, Location, Location: Compartmentalization of NAD+ Synthesis and Functions in Mammalian Cells. Trends in Biochemical Sciences 45:858–873. DOI: 10.1016/j.tibs.2020.05.010, PMID: 32595066

Covarrubias AJ, Perrone R, Grozio A, Verdin E. 2021. NAD+ metabolism and its roles in cellular processes during ageing. Nature reviews. Molecular cell biology 22:119–141. DOI: 10.1038/s41580-020-00313-x, PMID: 33353981

D’Alessandro KB, Zampese E, Blum JLE, Kuusik B, Palmiotti A, Davidson SM, Reczek CR, Surmeier DJ, Chandel NS. 2025. Genetic modulation of mitochondrial NAD+ regeneration does not prevent dopaminergic neuron dysfunction caused by mitochondrial complex I impairment. Frontiers in Cell and Developmental Biology 13:1650462. DOI: 10.3389/fcell.2025.1650462, PMID: 41081041

Diebold LP, Gil HJ, Gao P, Martinez CA, Weinberg SE, Chandel NS. 2019. Mitochondrial complex III is necessary for endothelial cell proliferation during angiogenesis. Nature metabolism 1:158–171. DOI: 10.1038/s42255-018-0011-x, PMID: 31106291

Duncan OF, Granat L, Ranganathan R, Singh VK, Mazaud D, Fanto M, Chambers D, Ballard CG, Bateman JM. 2018. Ras-ERK-ETS inhibition alleviates neuronal mitochondrial dysfunction by reprogramming mitochondrial retrograde signaling. PLoS Genetics 14:e1007567. DOI: 10.1371/journal.pgen.1007567, PMID: 30059502

Erdem A, Kaye S, Caligiore F, Johanns M, Leguay F, Schuringa JJ, Ito K, Bommer G, van Gastel N. 2025. Lactate dehydrogenase A-coupled NAD+ regeneration is critical for acute myeloid leukemia cell survival. Cancer & Metabolism 13:22. DOI: 10.1186/s40170-025-00392-4

Freije WA, Mandal S, Banerjee U. 2012. Expression Profiling of Attenuated Mitochondrial Function Identifies Retrograde Signals in Drosophila. G3: Genes|Genomes|Genetics 2:843–851. DOI: 10.1534/g3.112.002584, PMID: 22908033

Garcia S, Saldana-Caboverde A, Anwar M, Raval AP, Nissanka N, Pinto M, Moraes CT, Diaz F. 2022. Enhanced glycolysis and GSK3 inactivation promote brain metabolic adaptations following neuronal mitochondrial stress. Human Molecular Genetics 31:692–704. DOI: 10.1093/hmg/ddab282, PMID: 34559217

Granat L, Knorr DY, Ranson DC, Hamer EL, Chakrabarty RP, Mattedi F, Fort-Aznar L, Hirth F, Sweeney ST, Vagnoni A, Chandel NS, Bateman JM. 2023. Yeast NDI1 reconfigures neuronal metabolism and prevents the unfolded protein response in mitochondrial complex I deficiency. PLoS genetics 19:e1010793. DOI: 10.1371/journal.pgen.1010793, PMID: 37399212

Gray LR, Tompkins SC, Taylor EB. 2013. Regulation of pyruvate metabolism and human disease. Cellular and Molecular Life Sciences: CMLS 71:2577–2604. DOI: 10.1007/s00018-013-1539-2, PMID: 24363178

Hunt RJ, Granat L, McElroy GS, Ranganathan R, Chandel NS, Bateman JM. 2019. Mitochondrial stress causes neuronal dysfunction via an ATF4-dependent increase in L-2-hydroxyglutarate. The Journal of Cell Biology 218:4007–4016. DOI: 10.1083/jcb.201904148, PMID: 31645461

Jiménez-Gómez B, Ortega-Sáenz P, Gao L, González-Rodríguez P, García-Flores P, Chandel N, López-Barneo J. 2023. Transgenic NADH dehydrogenase restores oxygen regulation of breathing in mitochondrial complex I-deficient mice. Nature Communications 14:1172. DOI: 10.1038/s41467-023-36894-2, PMID: 36859533

Kanamori Y, Saito A, Hagiwara-Komoda Y, Tanaka D, Mitsumasu K, Kikuta S, Watanabe M, Cornette R, Kikawada T, Okuda T. 2010. The trehalose transporter 1 gene sequence is conserved in insects and encodes proteins with different kinetic properties involved in trehalose import into peripheral tissues. Insect Biochemistry and Molecular Biology 40:30–37. DOI: 10.1016/j.ibmb.2009.12.006, PMID: 20035867

Kang EY-C, Tseng Y-J, Peng W-H, Hung H-C, Lin P-H, Montales KP, Sherman E, Peregrin J, Wang EH, Kang C, Teng Y-C, Huang C-Y, Tsai C-L, Chang IY-F, Chen J, Tezel G, He Y, Li T-D, Stiles L, Shirihai O, Tsang SH, Lai C-C, Tsai C-N, Lin C-S, Wang N-K. 2026. Disrupted energy metabolism is associated with retinal ganglion cell degeneration in autosomal dominant optic atrophy. Science Advances 12:eadx7815. DOI: 10.1126/sciadv.adx7815, PMID: 41706861

Köhler-Solís A, Schirmeier S. 2025. Neuronal metabolism: Surprisingly flexible. Current Biology 35:R1098–R1101. DOI: 10.1016/j.cub.2025.10.010, PMID: 41253123

Kolde R. 2019. pheatmap: Pretty Heatmaps_. R package version 1.0.12.

Larsson NG, Wang J, Wilhelmsson H, Oldfors A, Rustin P, Lewandoski M, Barsh GS, Clayton DA. 1998. Mitochondrial transcription factor A is necessary for mtDNA maintenance and embryogenesis in mice. Nature Genetics 18:231–236. DOI: 10.1038/ng0398-231, PMID: 9500544

Long DM, Frame AK, Reardon PN, Cumming RC, Hendrix DA, Kretzschmar D, Giebultowicz JM. 2020. Lactate dehydrogenase expression modulates longevity and neurodegeneration in Drosophila melanogaster. Aging (Albany NY) 12:10041–10058. DOI: 10.18632/aging.103373, PMID: 32484787

Love MI, Huber W, Anders S. 2014. Moderated estimation of fold change and dispersion for RNA-seq data with DESeq2. Genome Biology 15:550. DOI: 10.1186/s13059-014-0550-8

Luengo A, Li Z, Gui DY, Sullivan LB, Zagorulya M, Do BT, Ferreira R, Naamati A, Ali A, Lewis CA, Thomas CJ, Spranger S, Matheson NJ, Vander Heiden MG. 2021. Increased demand for NAD+ relative to ATP drives aerobic glycolysis. Molecular cell 81:691–707.e6. DOI: 10.1016/j.molcel.2020.12.012, PMID: 33382985

Magistretti PJ, Allaman I. 2015. A Cellular Perspective on Brain Energy Metabolism and Functional Imaging. Neuron 86:883–901. DOI: 10.1016/j.neuron.2015.03.035, PMID: 25996133

Magrassi L, Leto K, Rossi F. 2013. Lifespan of neurons is uncoupled from organismal lifespan. Proceedings of the National Academy of Sciences of the United States of America 110:4374–4379. DOI: 10.1073/pnas.1217505110, PMID: 23440189

McElroy GregoryS, Reczek CR, Reyfman PA, Mithal DS, Horbinski CM, Chandel NS. 2020. NAD+ regeneration rescues lifespan but not ataxia in a mouse model of brain mitochondrial complex I dysfunction. Cell metabolism 32:301–308.e6. DOI: 10.1016/j.cmet.2020.06.003, PMID: 32574562

McMullen E, Weiler A, Becker HM, Schirmeier S. 2021. Plasticity of Carbohydrate Transport at the Blood-Brain Barrier. Frontiers in Behavioral Neuroscience 14:612430. DOI: 10.3389/fnbeh.2020.612430, PMID: 33551766

Mick E, Titov DV, Skinner OS, Sharma R, Jourdain AA, Mootha VK. 2020. Distinct mitochondrial defects trigger the integrated stress response depending on the metabolic state of the cell. eLife 9:e49178. DOI: 10.7554/eLife.49178

Motori E, Atanassov I, Kochan SMV, Folz-Donahue K, Sakthivelu V, Giavalisco P, Toni N, Puyal J, Larsson N-G. 2020. Neuronal metabolic rewiring promotes resilience to neurodegeneration caused by mitochondrial dysfunction. Science Advances 6:eaba8271. DOI: 10.1126/sciadv.aba8271, PMID: 32923630

Murali Mahadevan H, Hashemiaghdam A, Ashrafi G, Harbauer AB. 2021. Mitochondria in Neuronal Health: From Energy Metabolism to Parkinson’s Disease. Advanced Biology 5:e2100663. DOI: 10.1002/adbi.202100663, PMID: 34382382

Perkins LA, Holderbaum L, Tao R, Hu Y, Sopko R, McCall K, Yang-Zhou D, Flockhart I, Binari R, Shim H-S, Miller A, Housden A, Foos M, Randkelv S, Kelley C, Namgyal P, Villalta C, Liu L-P, Jiang X, Huan-Huan Q, Wang X, Fujiyama A, Toyoda A, Ayers K, Blum A, Czech B, Neumuller R, Yan D, Cavallaro A, Hibbard K, Hall D, Cooley L, Hannon GJ, Lehmann R, Parks A, Mohr SE, Ueda R, Kondo S, Ni J-Q, Perrimon N. 2015. The Transgenic RNAi Project at Harvard Medical School: Resources and Validation. Genetics 201:843–852. DOI: 10.1534/genetics.115.180208

Price MS, Rastegari E, Gupta R, Vo K, Moore TI, Venkatachalam K. 2025. Intracellular lactate dynamics in Drosophila neurons. iScience 28:113462. DOI: 10.1016/j.isci.2025.113462, PMID: 41169509

Quintana A, Kruse SE, Kapur RP, Sanz E, Palmiter RD. 2010. Complex I deficiency due to loss of Ndufs4 in the brain results in progressive encephalopathy resembling Leigh syndrome. Proceedings of the National Academy of Sciences of the United States of America 107:10996–11001. DOI: 10.1073/pnas.1006214107, PMID: 20534480

Rafikov R, Sun X, Rafikova O, Louise Meadows M, Desai AA, Khalpey Z, Yuan JX-J, Fineman JR, Black SM. 2015. Complex I dysfunction underlies the glycolytic switch in pulmonary hypertensive smooth muscle cells. Redox Biology 6:278–286. DOI: 10.1016/j.redox.2015.07.016, PMID: 26298201

Requejo-Aguilar R, Lopez-Fabuel I, Fernandez E, Martins LM, Almeida A, Bolaños JP. 2014. PINK1 deficiency sustains cell proliferation by reprogramming glucose metabolism through HIF1. Nature Communications 5:4514. DOI: 10.1038/ncomms5514

Richhariya S, Shin D, Le JQ, Rosbash M. 2023. Dissecting neuron-specific functions of circadian genes using modified cell-specific CRISPR approaches. Proceedings of the National Academy of Sciences 120:e2303779120. DOI: 10.1073/pnas.2303779120

Richhariya S, Shin D, Schlichting M, Rosbash M. 2025. Metabolic rewiring prevents neurodegeneration caused by chronic mitochondrial dysfunction. Current Biology 35:5443–5459.e5. DOI: 10.1016/j.cub.2025.09.063, PMID: 41151583

Rodrigues APC, Novaes AC, Ciesielski GL, Oliveira MT. 2022. Mitochondrial DNA maintenance in Drosophila melanogaster. Bioscience Reports 42:BSR20211693. DOI: 10.1042/BSR20211693, PMID: 36254835

Sanz A, Soikkeli M, Portero-Otín M, Wilson A, Kemppainen E, McIlroy G, Ellilä S, Kemppainen KK, Tuomela T, Lakanmaa M, Kiviranta E, Stefanatos R, Dufour E, Hutz B, Naudí A, Jové M, Zeb A, Vartiainen S, Matsuno-Yagi A, Yagi T, Rustin P, Pamplona R, Jacobs HT. 2010. Expression of the yeast NADH dehydrogenase Ndi1 in Drosophila confers increased lifespan independently of dietary restriction. Proceedings of the National Academy of Sciences of the United States of America 107:9105–9110. 10.1073/pnas.0911539107, PMID: 20435911

Schindelin J, Arganda-Carreras I, Frise E, Kaynig V, Longair M, Pietzsch T, Preibisch S, Rueden C, Saalfeld S, Schmid B, Tinevez J-Y, White DJ, Hartenstein V, Eliceiri K, Tomancak P, Cardona A. 2012. Fiji: an open-source platform for biological-image analysis. Nature Methods 9:676–682. DOI: 10.1038/nmeth.2019

Schmidt O, Pfanner N, Meisinger C. 2010. Mitochondrial protein import: from proteomics to functional mechanisms. Nature Reviews Molecular Cell Biology 11:655–667. DOI: 10.1038/nrm2959

Sorge S, Theelke J, Yildirim K, Hertenstein H, McMullen E, Müller S, Altbürger C, Schirmeier S, Lohmann I. 2020. ATF4-Induced Warburg Metabolism Drives Over-Proliferation in Drosophila. Cell Reports 31:107659. DOI: 10.1016/j.celrep.2020.107659, PMID: 32433968

Supplie LM, Düking T, Campbell G, Diaz F, Moraes CT, Götz M, Hamprecht B, Boretius S, Mahad D, Nave K-A. 2017. Respiration-Deficient Astrocytes Survive As Glycolytic Cells In Vivo. The Journal of Neuroscience: The Official Journal of the Society for Neuroscience 37:4231–4242. DOI: 10.1523/JNEUROSCI.0756-16.2017, PMID: 28314814

Titov DV, Cracan V, Goodman RP, Peng J, Grabarek Z, Mootha VK. 2016. Complementation of mitochondrial electron transport chain by manipulation of the NAD+/NADH ratio. Science 352:231–235. DOI: 10.1126/science.aad4017, PMID: 27124460

Vecsey CG, Koochagian C, Porter MT, Roman G, Sitaraman D. 2024. Analysis of Sleep and Circadian Rhythms from Drosophila Activity-Monitoring Data Using SCAMP. Cold Spring Harbor protocols 2024:pdb.prot108182. DOI: 10.1101/pdb.prot108182, PMID: 38336392

Veits N, Guo Y, He J, Mazouni K, Nemazanyy I, Bres M, Picciotto C, Mestdagh C, Yan Y, Schweisguth F. 2026. NAD+ supply and redox state limit developmental speed in the Drosophila eye. The EMBO Journal 45:4094–4123. DOI: 10.1038/s44318-026-00801-4

Vercellino I, Sazanov LA. 2022. The assembly, regulation and function of the mitochondrial respiratory chain. Nature Reviews Molecular Cell Biology 23:141–161. DOI: 10.1038/s41580-021-00415-0

Volkenhoff A, Weiler A, Letzel M, Stehling M, Klämbt C, Schirmeier S. 2015. Glial Glycolysis Is Essential for Neuronal Survival in Drosophila. Cell Metabolism 22:437–447. DOI: 10.1016/j.cmet.2015.07.006, PMID: 26235423

Yadav S, Pan X, Li S, Martin PL, Hoang N, Chen K, Karhadkar A, Malhotra J, Zuckerman AL, Munan S, Klose MK, Wang L, Cracan V, Parkhitko AA. 2026. Perturbation of NAD(P)H metabolism with the LbNOX xenotopic tool extends lifespan and mitigates age-related changes. Science Advances 12:eady0628. DOI: 10.1126/sciadv.ady0628

